# Pre-meiotic H1.1 degradation is essential for Arabidopsis gametogenesis

**DOI:** 10.1101/2025.04.18.649492

**Authors:** Yanru Li, Danli Fei, Jasmin Schubert, Kinga Rutowicz, Zuzanna Kaczmarska, Alberto Linares, Alejandro Giraldo Fonseca, Sylvain Bischof, Ueli Grossniklaus, Célia Baroux

## Abstract

Despite being evolutionary distant, plants and animals share a common phenomenon at the somatic-to-reproductive cell fate transition with extensive structural and compositional changes in chromatin. Such chromatin reprogramming occurs in the plant SMCs (Spore Mother Cells) and animal PGCs (primordial germ cells) and is initiated by the loss of linker histones (H1). H1 loss is essential to establish pluripotency in animal PGCs but its role is not known in plants. Here, we identified two regulatory pathways involving a citrullinase and an E3-ubiquitin ligase that contribute H1.1 loss in female SMC in Arabidopsis. We also identified roles for two specific residues: an arginine, whose positive charge contributes to H1.1 destabilization from chromatin, and a lysine in the globular domain that is essential for H1.1 degradation. Ovules with impaired H1.1 loss in the SMC proceed through sporogenesis but fail to complete gametogenesis. We propose a citrullination-ubiquitination pathway governing pre-meiotic H1 depletion as a critical mechanism for establishing post-meiotic competence in the *Arabidopsis* germline.

## INTRODUCTION

Sexually reproducing multicellular organisms form a specialized cell lineage dedicated to meiosis and gametogenesis, two processes that are fundamental to ensuring genetic diversity while maintaining chromosomal integrity across generations. In mammals, this lineage arises from primordial germ cells (PGCs), which are specified early in development and set aside from the pluripotent embryonic cell mass. Following meiosis, these cells directly differentiate into specialized gametes (Bendel–Stenzel *et al*, 1998). In flowering plants, the functional equivalent of PGCs are spore mother cells (SMCs), which form later in development within specific floral tissues of the adult organism. In contrast to animal meiotic products, the haploid spores of plants are pluripotent and give rise to a multicellular gametophyte, containing several distinct cell types, including the gametes (Skinner & Sundaresan, 2018). This difference in timing, origin, and developmental competence of the meiotic products raises questions about the mechanisms that regulate the *de novo* specification of SMCs in plants and the establishment of a pluripotent state following meiosis.

Recent studies have uncovered multiple layers of regulation controlling the establishment of female SMC in the model plant *Arabidopsis*. Their specification involves geometrical information, hormonal cues, cell-cell signaling modules, genetic and epigenetic factors (Pinto *et al*, 2019; Lora *et al*, 2019; Jiang & Zheng, 2022; Hernandez-Lagana *et al*, 2021; Cai *et al*, 2022, 2025; Huang *et al*, 2023). Typically, female SMCs are initiated by the enlargement of the most distal cell in the subepidermal layer (L2) of ovule primordia (Pinto *et al*, 2019; Lora *et al*, 2019). SMC differentiation, defined by distinct morphological features and markers, is a gradual process tightly linked to the growth of the ovule primordium(Hernandez-Lagana *et al*, 2021). While the mechanisms are not fully elucidated, several non-cell autonomous mechanisms have been identified that govern both SMC formation and singleness. SMC specification involves a transcriptional co-repressor complex, consisting of SPOROCYTELESS/NOZZLE (SPL/NZZ) (Balasubramanian & Schneitz, 2000) and TOPLESS/TOPLESS-RELATED (TPL/TPR) proteins (Wei *et al*, 2015). This complex likely acts indirectly to enable the expression of *WUSCHEL* (WUS), a homeodomain transcription factor. WUS subsequently activates the expression of *WINDHOSE1* (WIH1) and *WIH2* at the apex of the primordium that, together with the tetraspanin-type protein *TORNADO2* (TRN2)/EKEKO, contribute to initiate SMC specification (Lieber *et al*, 2011; Pinto *et al*, 2019; Lora *et al*, 2019). Furthermore, the singleness of SMCs is controlled by lateral inhibition involving small interfering RNAs (siRNAs) and hormones, i.e. auxin and brassinosteroids, that prevent neighboring cells from adopting an SMC fate (Cai *et al*, 2022; Lora *et al*, 2019; Grossniklaus & Schneitz, 1998; Cai *et al*, 2025). Growth of the primordium itself also contributes to SMC singleness because the shape of its apex likely constrain the domain of action of key regulators together with mechanical cues supporting SMC expansion (Hernandez-Lagana *et al*, 2021). SMC singleness is further secured by cell cycle regulators, including KIP-RELATED PROTEINS (KRPs), also known as INHIBITORS OF CYCLIN-DEPENDENT KINASES (ICKs), and CYCLIN-DEPENDENT KINASES (CDKs), which help prevent mitotic division of the SMC, by inhibiting RETINOBLASTOMA-RELATED1 (RBR1), an inhibitor of WUS (Zhao *et al*, 2017; Cao *et al*, 2018). Thus, SMC specification results from the spatio-temporal integration of an array of intrinsic and extrinsic signals.

Ultimately, SMCs are committed to undergo meiosis, producing functional haploid spores that give rise to gametophytes that harbor the gametes. The SMC transcriptome exhibits signatures of reprogramming consistent with cell fate change (Schmidt *et al*, 2011; Hou *et al*, 2021) but the underlying epigenome has largely remained inaccessible to profiling studies. Yet, the SMC chromatin undergoes drastic structural and compositional changes compared to that of neighboring cells. These changes include the loss of linker histone variants H1.1 and H1.2 and of nucleosomal histone variants H3.1/HTR13 and H2AZ/HTA11, elevated levels of H3K4me3 and H4ac, reduced levels of H3K27me3 and DNA methylation in the CHH context (Ingouff *et al*, 2017; She *et al*, 2013). Interestingly, however, the overall transcriptional activity remains low in SMCs compared to neighboring cells (She *et al*, 2013). The loss of H1 linker histones in Arabidopsis SMCs (She *et al*, 2013) is reminiscent of the loss of somatic H1 subtypes (H1.1–H1.5 and H1.10) in mouse PGCs during early pre-implantation stage (E11.5) (Hajkova *et al*, 2008; Izzo *et al*, 2017). H1 variants were long considered to serve only a structural role in chromatin folding hindering transcription. But recent studies uncovered a more complex interaction between H1 histones and epigenetic regulation, including DNA methylation, as well as, histone methylation, and acetylation, both in plants and animals (Bendel–Stenzel *et al*, 1998; Skinner & Sundaresan, 2018; Koltunow & Grossniklaus, 2003; Pinto *et al*, 2019; Lora *et al*, 2019; Jiang & Zheng, 2022; Hernandez-Lagana *et al*, 2021). This raises the question whether the loss of H1 in plant SMCs and mouse PGCs controls cellular reprogramming associated with the somatic-to-reproductive transition. Supporting this hypothesis, H1 depletion in mouse PGC was found critical for establishing pluripotency within the germline (Christophorou *et al*, 2014). Similarly, H1 depletion in mouse embryonic stem cells leads to the upregulation of pluripotency genes, causing the cells to stall in a self-renewal state with impaired differentiation potential (Zhang *et al*, 2012).

In plants, the role and mechanisms of H1 loss in Arabidopsis SMCs are unknown. Here we report on two pathways involving the citrullinase AGMATINE IMINO HYDROLASE (AIH) and the E3 ubiquitin ligase CULLIN4 that regulate H1.1 loss in the SMC, involving two key residues of H1.1, an arginine R57 and lysin K89. We propose a working model involving arginine citrullination as a mechanism increasing H1.1 mobility and ubiquitination leading to degradation. Importantly, we found that disrupting H1 loss in the SMC does not affect meiosis but instead compromises gametogenesis suggesting that pre-meiotic chromatin dynamics contributes to post-meiotic fate.

## RESULTS

### The E3-ubiquitin ligase CULLIN4 contributes to H1.1 degradation in the SMC

In previous work, we reported the loss of H1.1 and H1.2 in *Arabidopsis* male and female SMCs but not in neighboring cells (She & Baroux, 2015, She, Grimanelli et al., 2013). As this process was sensitive to the proteasome inhibitor Syringolin A(She *et al*, 2013) we hypothesized a ubiquitin-mediated targeting of H1 to the proteasome, a major pathway for protein degradation in plants (Vierstra, 1993). E3 ligases of the CULLIN family which transfer ubiquitin to the target protein (Yang *et al*, 2021) were thus prime candidates. Published RNA *in situ* hybridization indicated *CULLIN4 (CUL4)* expression in young ovule primordia (Chen *et al*, 2006) and microarray data confirmed expression in female SMCs and surrounding nucellus (**Figure EV1A**). Thus, if CULLIN4 were involved in H1 degradation, its downregulation at the onset of ovule primordium development should prevent loss of in mature SMCs. To avoid secondary effects arising from constitutive loss-of-function mutations, we conditionally knocked-down *CUL4* in developing flower buds. To this aim, we expressed an artificial microRNA targeting *CUL4, amiR[CUL4]* (**Figure EV1B**) under control of the *pOP/LhGR* Dexamethasone (Dex)-inducible system (Craft *et al*, 2005; Samalova *et al*, 2019). We confirmed a reduction of CUL4 protein levels in seedlings induced for *amiR[CUL4]* expression and observed the known *fusca* phenotype typical of a *cul4* loss-of-function mutation (Chen *et al*, 2006) (**Figure EV1C, D**). To investigate the possible role of CUL4 in H1 depletion during SMC development, we introgressed the inducible *amiR[CUL4]* line into an H1.1-GFP reporter line (She et al., 2013). We rationalized that if if CUL4 played a role in this process, knock-down of its expression should lead to partial or full retention of H1.1-GFP in the SMC. *amiR[CUL4]* expression was induced in young flower buds *in planta* as previously described (Schubert, Li et al., 2022). Five days post induction (5 dpi), flower buds were collected, and ovule primordia selectively at stage 2-I were dissected and scored by confocal microsocopy for two categories: those showing no detectable H1.1 in the SMC (“depletion”) and those showing a detectable signal (“persistence”). The Mock control showed 11% (n=137) primordia with a residual H1.1-GFP signal, likely due to the gradual nature of H1 loss. By contrast, Dex-treated flower buds harbored 52% (n=178) primordia with a strong H1.1-GFP signal in the SMC (**Figure 1A**). We obtained similar results when primordia were treated with the proteasome inhibitor epoxomicin (**Figure 1B**). We concluded that *amiR[CUL4]* expression in young ovule primordia can impair H1.1 depletion during SMC formation, likely due to effective *CUL4* downregulation. The delicate experimental setup, involving manual treatment of young flower buds for induction and the use of an artificial miRNA likely explains why H1 persistence was not fully penetrant. But this approach has the benefit of allowing spatial and temporal control minimizing confounding effects caused by constitutive loss-of-function. Collectively, our results indicate that the E3-ubiquitin ligase CULLIN4 plays a role in H1.1 degradation during SMC development.

**Figure 1.**
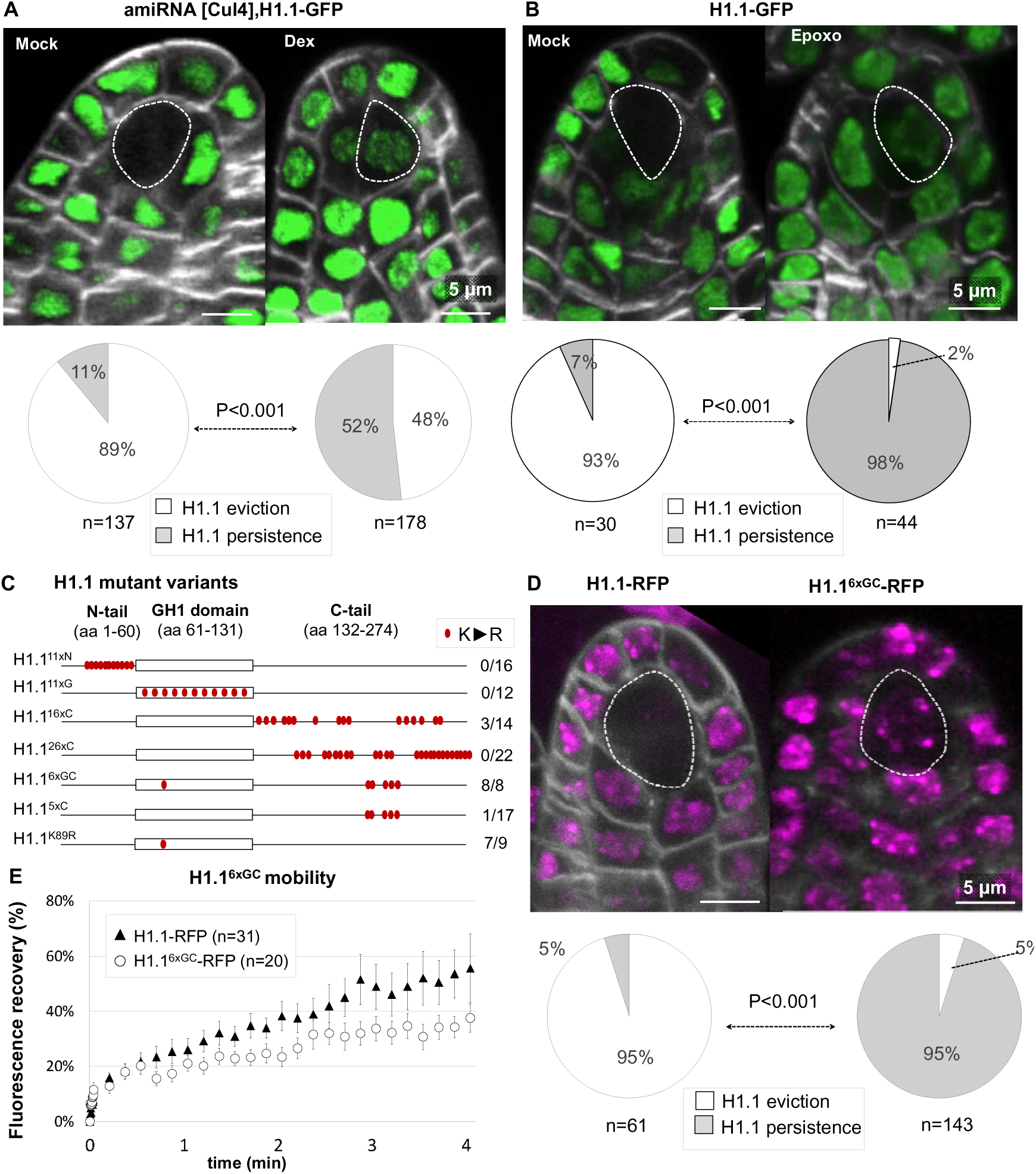
H1.1 degradation is mediated by the E3-Ubiquitin ligase CULLIN 4. **(A-B)** H1.1-GFP degradation in the SMC is impaired when inducing the expression of an artificial miRNA against *CUL4* (**A,** *amiRNA[CUL4]*) or when treating with the proteasome inhibitor epoxomicin (**B**). Young inflorescences were treated either with 10µM dexamethasone or a mock solution as control (A, Dex, Mock) (Craft *et al*, 2005; Samalova *et al*, 2019), or with a 5µM epoxomicin solution or mock solution as control (B, Epoxo, Mock). Images shows partial projections of confocal images showing the GFP signal (green) and cell wall stained with Renaissance (Musielak *et al*, 2015) (grey). The SMC is indicated with dotted contours. Pie charts below the images show the proportion of ovule primordia at stage 2-I/2-II 5 days post induction (5 dpi) showing either H1.1-GFP depletion (white) or persistence (grey) in the SMC. n, number of primordia scored. P value from a Fischer-exact test. Replicate experiments in Supplemental data 1, scoring data in Data Source 1. **(C)** Schematic representation of H1.1 mutant variants carrying several K-to-R amino acid substitutions (red dots) in their N-terminal tail, globular domain, C-terminal tail, or a combination thereof, and fused to RFP. The numbers on the right indicate the number of independent lines showing a persistent signal in the SMC, indicating a lack of degradation, over (/) the total number of lines screened. (**D**) Representative images of H1.1-RFP and H1.1^6xGC^RFP expression in ovule primordia at stage 2-I, 5 dpi. Pie charts show the proportion of ovule primordia showing either H1.1-RFP depletion (white) or persistence (grey) in the SMC. n, number of primordia scored. P value from a Fischer-exact test. Replicate experiments in Supplemental data 1, scoring data in Data Source 1. **(E)** Fluorescence recovery after photobleaching (FRAP) of H1.1-RFP and H1.1^6xGC^RFP in root nuclei from 5 days old seedlings grown in Dex. n, number of analyzed nuclei. Statistical analysis in Figure EV1, raw data in Data Source 1.

### A screen for potential ubiquitination targets identifies K89 in the globular domain of H1.1

E3-ligases preferentially target lysine-rich regions (Yang *et al*, 2021). Given that linker histones are particularly rich in lysine throughout their protein sequence (Jerzmanowski, 2004), we considered that ubiquitination of one or several lysines could drive H1 degradation. To test this hypothesis, we first created six H1.1 mutant variants with batches of K-to-R amino acid substitutions (**Figure 1C**). Candidate lysine were selected based on computational predictions (Chen *et al*, 2011; Radivojac *et al*, 2010; Walton *et al*, 2016), proteomic evidence of ubiquitination in seedling tissues (Walton *et al*, 2016) and comparison with ubiquitinated sites in mouse and human counterparts (Wiśniewski *et al*, 2007). These mutant H1.1 variants were designed to carry K-to-R mutations in the N-terminal domain, the Globular domain, the C-terminal tail or a combination thereof. The mutant variants were expressed as RFP translational fusions under the control of the Dex-inducible *pOP/LhGR* system as before (Craft *et al*, 2005; Samalova *et al*, 2019). We screened primary transformants following induction on young flower buds for H1.1-RFP depletion versus persistence in the SMC of ovule primordia at stage 2-I/2-II by confocal microscopy. None of the 16 and 12 transformants expressing mutant variants carrying 11 K-to-R substitution in the N-terminal tail (H1.1^11xN^) or the globular domain (H1.1^11xG^), respectively, showed persistence either did any of the 26 transformants expressing a mutant variant with 26 substitutions in the C-terminal tail (H1.1^26xC^), (**Figure 1C**). In 3 lines expressing a variant with 16 substitutions in the C-terminal tail (H1.1^16xC^) we observed some residual H1.1^16xC^-RFP signal but since it was not reproduced in the other 11 independent transformants we did not consider this mutant variant further (**Figure 1C**). However, all of the eight transformants expressing H1.1^6xGC^, a variant with 6 substitutions (five in the C-terminal tail, one in the globular domain) showed a strong persistence of H1.1^6xGC^-RFP in the SMC (**Figure 1C, D**). This H1.1 variant was mutated in all six sites shown to be ubiquitinated in seedling (Walton *et al*, 2016), (**Figure EV1E**). Typically, 76-95% of the ovule primordia expressing H1.1^6xGC^-RFP showed a clear signal in the SMC at stage 2-I / 2-II (**Figure 1D** and **Figure S1F** for replicates). In contrast, primordia expressing a native H1.1 variant fused to RFP (H1.1-RFP) under control of the same Dex-inducible system, showed depletion in 95% of the SMCs in ovule primordia at stage 2-I/2-II (**Figure 1D**), with the remaining 5% showing only weak, residual signals. The persistence of H1.1^6xGC^-RFP could be due to either a lack of ubiquitination and thus degradation, or enhanced stability on chromatin. Because profiling H1.1 post-translational modifications (PTMs) in Arabidopsis is currently out of reach we assessed H1.1^6xGC^ stability by Fluorescence Recovery After Photobleaching (FRAP). FRAP was previously used to investigate turnover and residency time of H1 mutant variants *in vivo*, in order to identify residues involved in binding the nucleosome (Brown *et al*, 2006).We compared the fluorescence recovery rates of H1.1-RFP and H1.1^6xGC^-RFP following photobleaching in Dex-treated seedling root nuclei. The data indicated a similar mobility, both during the initial, rapid recovery phase and later, during the slow recovery phase (**Figure 1E** and **Figure EV1G**). This suggests that substitution of the six lysines into arginines – harboring a similar charge-did not enhance H1.1 stability. Considering that these lysines are ubiquitinated in seedling tissues (Walton *et al*, 2016), our findings strongly suggest that ubiquitination, and consequently, degradation of H1.1^6xGC^-RFP is compromised.

To further distinguish the contribution of the different lysine mutated in the H1.1^6xGC^ variant, we created two additional mutants: one carrying a single mutation at lysine 89 (K89R) in the globular domain (H1.1^K89R^, **Figure 1C**, **Figure EV1H**) and another with K-to-R mutations at the five remaining lysines in the C-terminal tail (H1.1^5xC^, **Figure 1C**). We expressed these variants under the control of the Dex-inducible system and screened several independent transformants. The persistence phenotype was recapitulated in 7 of 9 lines expressing H1.1^K89R^-RFP but in none of 16 lines expressing H1.1^5xC^-RFP indicating that a single amin acid change is sufficient to render H1.1^K89R^-RFP resistant to degradation. K89 is exposed at the surface of the third alpha-helix of the globular domain (**Figure EV1H**); hence, is unlikely to contribute to interactions with the nucleosome. In line with this idea, mutation of K52 in the mouse MmH1(o) variant, homologous to K89 in AtH1.1 (**Figure EV1H**) did not change its mobility(Brown *et al*, 2006). Similarly, the K89R mutation in the *Arabidopsis* variant did not affect the overall mobility of H1.1 (**Figure EV1G**). Finally, we observed that both H1.1^6xGC^-RFP and H1.1^K89R^-RFP showed a nuclear distribution like the native H1.1-RFP, present in both euchromatin and heterochromatin (**Figure EV1J**).

In conclusion, we identified a conserved lysine residue, K89, in the globular domain of H1.1 that plays a key role in the degradation of H1.1 in the SMC. The fact that this lysine does not affect binding affinity and was found to be ubiquitinated in seedlings strongly supports a model in which ubiquitination at K89, likely mediated by CUL4, directly or indirectly, drives H1.1 degradation.

### A single arginine residue in the N-terminal tail controls H1 depletion in the SMC

Ubiquitin-mediated protein degradation is a basic cellular process and CUL4 is expressed throughout the ovule primordium (Dumbliauskas *et al*, 2011). Hence, we wondered whether another PTM of H1.1 might prime H1.1 for ubiquitination specifically in the SMC. Interestingly, H1.2 depletion in mouse PGCs (Hajkova *et al*, 2008) is controlled by citrullination of R52, a PTM converting arginine into citrulline (Christophorou *et al*, 2014). We thus wondered whether a similar citrullination mechanism might operate in the control of H1.1 depletion in *Arabidopsis* SMCs. We created H1.1 mutant variants either with a lysine substitution at R57 (R57K) that ensures charge conservation, or an alanine substitution (R57A) which conveys a neutral charge similar to that of citrulline (**Figure 2A**). We expressed these mutant variants fused to RFP under the control of the *pOP/LhGR Dex*-inducible system. We scored the fraction of primordia showing presence or absence of RFP signal in the SMC of ovule primordia stage 2-I/2-II at 5dpi. We observed that the R57K mutation compromised H1.1 depletion in a large fraction of ovule primordia (26%, n=218 **Figure 2B** and 26%, n=209 **Figure EV2A**), but not as efficiently as the previously assessed K89R mutation, suggesting that these two residues may act differently on H1.1 degradation. In contrast, the H1.1^R57A^-RFP showed a residual signal in only 7% SMC with (n=171) like the H1.1-RFP control (**Figure EV2A, Figure 1D**). This finding further indicated that R57 is unlikely the target of a biochemical modification (for instance methylation) but rather suggests a role for its ionic charge. In addition, we observed that the R57K mutation also compromised depletion in male SMCs, whereas the R57A mutation did not (**Figure EV2B**).

**Figure 2.**
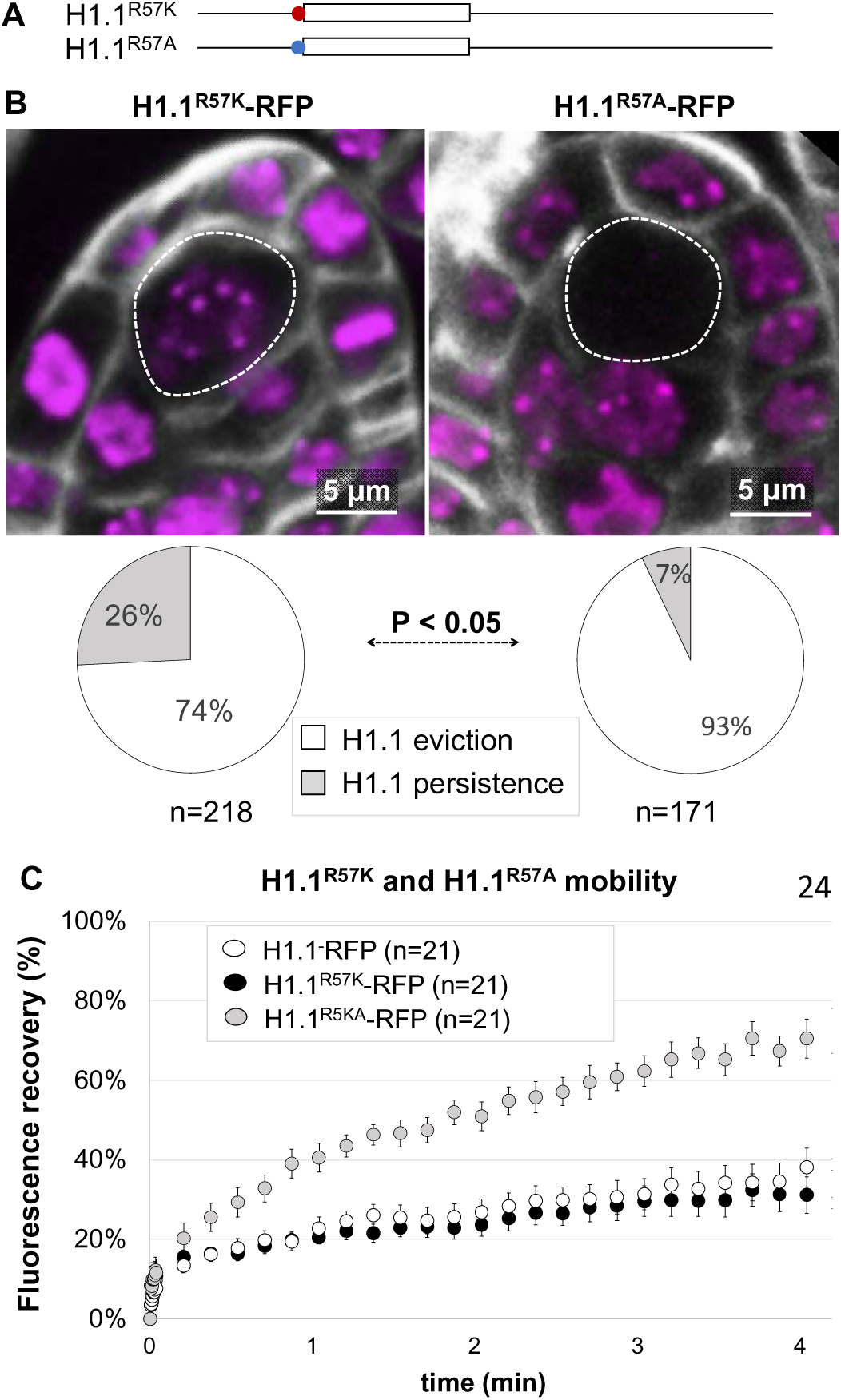
An R57K substitution prevents H1.1 depletion in SMC without affecting H1.1 mobility. **(A)** Schematic representation of the R57K and R57A mutation just before the globular domain (**B**) Representative images of H1.1^R57K^-RFP and H1.1^R57A^-RFP in ovule primordia stage 2-I at 5 dpi, showing persistence and depletion in the SMC (dotted lines), respectively (partial z-projections; RFP: magenta; Renaissance: grey). Pie charts below show the proportion of ovule primordia showing H1.1(mutant)-RFP depletion or persistence in the SMC. n, number of primordia scored. P value: Fischer exact test. Replicate experiments in Figure EV2. (**B)** Fluorescence recovery after photobleaching (FRAP) of H1.1^−^ RFP, H1.1^R57K^-RFP and H1.1^R57A^-RFP in root nuclei from 5 days old, Dex-induced seedlings. n, nuclei number. See Figure EV2 and Data Source 2.

To assess whether persistence of H1.1^R57K^-RFP in the SMC is due to enhanced chromatin stability compared to H1.1^R57A^-RFP or H1.1-RFP, we conducted FRAP analyses as previously described. Strikingly, the R57K mutation did not affect H1.1 mobility, whereas the R57A mutation significantly increased it, as evidenced by a two-fold increase recovery rate (**Figure 2C, Figure EV2C**). The stability of linker histones binding to the nucleosome is primarily governed by electrostatic interactions with positively charged amino acids on H1 (Brown *et al*, 2006; Martinsen *et al*, 2022). Hence, the most likely explanation for the reduced residency time - and faster recovery rate-of H1.1^R57A^ is that the alanine substitution causes charge neutralization and weaker electrostatic interactions between H1.1 and the nucleosome. In support of this hypothesis, we observed a more efficient depletion of H1.1^R57A^ in the SMC of ovule primordia after we refined the analysis by scoring primordia showing a faint signal in heterochromatin (“partial depletion”, **Figure EV2A**) which we interpret as an intermediate step to full depletion. We noticed that this fraction was significantly reduced in lines expressing H1.1^R57A^-RFP with a concomitant increase of SMC without signal (**Figure EV2A**), suggesting a more efficient depletion of H1.1^R57A^ than H1.1^R57K^ or H1.1.

### AGMATINASE IMINOHYDROLYASE, a candidate arginine deiminase is necessary for H1.1 depletion

Following up on our hypothesis that charge neutralization at R57 impacting H1.1 stability we investigated citrullination as a candidate mechanism. Citrullination driven by peptidyl arginine deiminases such as the PADI enzymes in animal cells, results in the net loss of a positive charge and has previously been shown to play a role in H1.2 depletion in mouse PGCs (Christophorou *et al*, 2014). The *Arabidopsis* genome does not encode proteins homologous to PADI enzymes. However, a 3D protein structure homology search identified AGMATINASE IMINOHYDROLYASE (AIH) as closely related to the catalytic domain of PADI4, with a striking conservation of the four catalytic amino acids (**Figure 3A**). AIH exhibits citrullinase activity *in vitro* (Marondedze *et al*, 2021) and is expressed in young flower buds at stage 8-10 according to publicly available transcriptome datasets (Klepikova *et al*, 2016) (**Figure EV3A**). Using RNA *in situ* hybridization, we confirmed that *AIH* is expressed in young ovule primordia (**Figure 3B**, **Figure EV3B**). To test whether AIH plays a role in H1.1 depletion we induced the expression of an artificial miRNA against *AIH* (*amiR[AIH]*, **Figure EV3C**) using the Dex-induction system, in an H1.1-GFP expressing line. Consistent with our hypothesis, H1.1-GFP depletion in the SMC was less efficient when *amiR[AIH]* was induced compared to mock-treated tissues, with 37% and 6% ovule primordia showing H1.1 persistence (n=30 and n=47), respectively (**Figure 3C** and **Figure EV3D**). To test the involvement of AIH independently, we used Cl-amidine, a known inhibitor of the mammalian citrullinase PADI4 that binds the catalytic domain(Luo *et al*, 2006), which is highly conserved in AIH (**Figure 3A**). Cl-amidine treatment also reduced H1.1 depletion in the SMC of a large fraction of ovule primordia (39%, n=56) compared to the mock treatment (**Figure 3D** and **Figure EV3E**).

**Figure 3.**
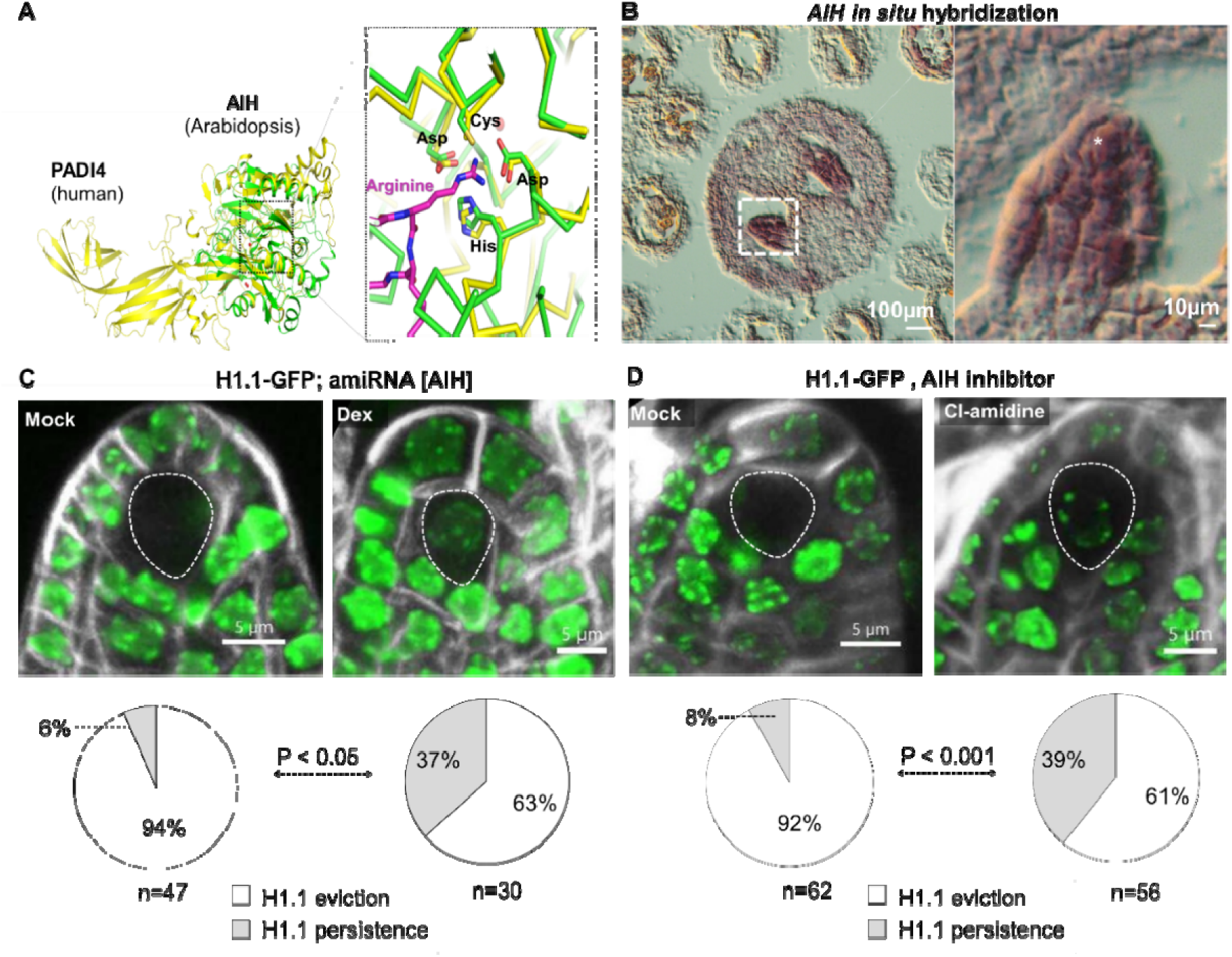
Citrullinase AIH Downregulation Impairs H1.1 Depletion in SMC. **(A)**.3D protein structure of the human citrullinase PADI4 (yellow, PDB model 2DEW) and the *Arabidopsis* citrullinase AIH (green, PDB model 1VKP) showing structural homology of AIH with the catalytic head of PADI4 and a perfect conservation of the catalytic site (Asp, Cys, Asp and His, inlet). **(B)**. Histochemical detection of *AIH* transcripts following RNA *in situ* hybridization on a transversal section of a flower bud stage 10 (left) revealing a specific signal in ovule primordia (box in dashed line and zoomed-in image on the right). **(C,D)** H1.1-GFP depletion in the SMC is prevented when inducing the expression of a downregulating artificial miRNA against *AIH* (*amiRNA[AIH]*, Dex versus mock, C) or when treating with the PADI4-specific enzyme inhibitor Cl-amidine (D). Images show partial projections of confocal images showing the GFP signal (green) and Renaissance cell boundary staining (grey). Pie charts below the images show the proportion of ovule primordia 5 dpi harboring no or persistent H1.1-GFP signal in the SMC. n, number of primordia scored. P value from a Fischer exact test. Replicate experiments in Figure EV3, Data Source 3.

In conclusion, genetic and toxicological evidence support a model in which AIH contributes to H1.1 depletion in the SMC.

### H1 persistence in the SMC affects chromatin reorganization without impairing SMC maturation or meiosis

Next, we asked whether compromising H1.1 depletion in the SMC may affect chromatin reorganization (She *et al*, 2013). We focused on key markers of constitutive and facultative heterochromatin, which decreases during the SMC chromatin reorganization (She *et al*, 2013; Ingouff *et al*, 2017): we measured the relative heterochromatin fraction (RHF) and the levels of H3K27me3 and methylated CHH readers, LHP1-GFP(Exner *et al*, 2009) and DynaMET mCHH-Venus (Ingouff *et al*, 2017), respectively. SMC showing with a persistent H1.1^R57K^-RFP signal showed higher levels of these three heterochromatin markers compared to SMCs without detectable levels of H1.1-RFP or H1.1^R57A^-RFP (**Figure 4A-C)**. By contrast, SMC expressing the persistent H1.1^6xGC^-RFP variant did not show significant changes for these markers compared to the control (**Figure C**, **Figure EV4A-B**). These findings suggest a different impact of the R57K and K89R substitutions on H1.1 properties relative to chromatin reorganization in the SMC.

**Figure 4.**
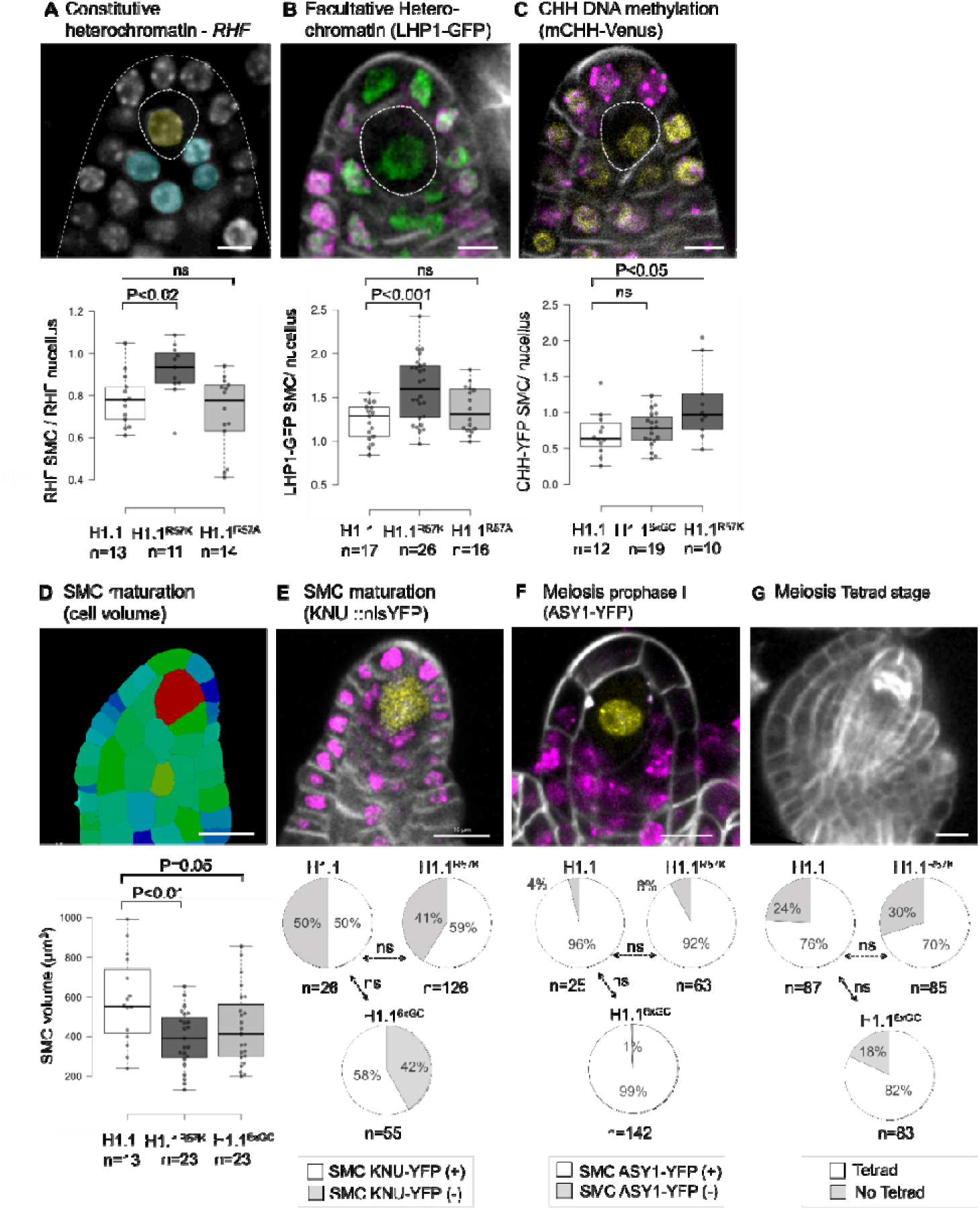
Inhibition of H1.1 depletion in the SMC disrupts chromatin reorganization without affecting meiosis. **(A-C)** Quantification of heterochromatin markers in the SMC of ovule primordia stage 2-I/2-II at 5 dpi in lines expressing H1.1-RFP, H1.1^R57K^-RFP, H1.1^R57A^-RFP or H1.1^6xGC^-RFP (H1.1, H1.1^R57K^, H1.1^R57A^, H1.1^6xGC^, respectively). Top: representative images (confocal projections) in H1.1-RFP control lines following whole-mount propidium iodide (PI) staining (A, yellow: SMC; blue: nucellus nuclei used as references); live GFP (B, green), YFP (C, yellow), RFP (B, C, magenta) and Renaissance counterstaining (B-C grey); Bottom: Graphs showing the relative heterochromatin fraction (RHF) in chromocenters (A), and relative LHP1-GFP (B) and mCHH-Venus (C) intensity in the SMC. The measurements are expressed relative to the nucellus. **(D-G)** Quantification of SMC maturation in ovule primordia stage 1-II/2-I at 5 dpi (D, E) and meiosis markers in ovule primordia stage 2-III/3-I at 7 dpi (F, G) in lines expressing H1.1-RFP, H1.1^R57K^-RFP, H1.1^R57A^-RFP, indicated above the pie charts as in A-C. Note that due to the progressive and variable expression of the *KNU::nlsYFP* marker, not all primordia showed a detectable signal also in the wild-type control. Top: representative images in the H1.1-RFP control line following: Renaissance staining, 3D imaging and cell segmentation with cells colored according to their volume (D), live imaging of *KNU::nlsYFP* (E, yellow), ASY1-YFP (F, yellow), H1.1-RFP (E-F, magenta), Renaissance counterstaining (E-G, grey) and confocal imaging (partial projection shown). Scale bars: 5µm (A-C), 10µm (D-G). P values, Mann-Whitney U test (A-D), Fisher exact test (E-G). n, number of ovule primordia. See also Figure EV4 and raw data in Data Source 4.

We then addressed whether persistence of H1.1 affects SMC differentiation and meiosis. We first analyzed SMC growth and the expression of the fate marker *KNU::nlsYFP*, two hallmarks of SMC maturation (Hernandez-Lagana & Autran, 2020; Tucker *et al*, 2003). We found that preventing H1.1 depletion by expressing either the H1.1^R57K^ or the H1.1^6GC^ mutant variant compromised SMC growth, as revealed by a 3D analysis of the primordium (**Figure 4D, Figure EVC**). In contrast, the size of companion cells increased (**Figure EVD**). Yet, *KNU::nlsYFP* expression was not altered in ovule primordia expressing either of the H1.1 mutant variant **(Figure 4E)**. Next, we assessed meiosis by scoring primordia expressing the ASY1-YFP marker, which is loaded onto chromatin in prophase I (Valuchova *et al*, 2020) and the occurrence of tetrads. Ovule primordia induced for the expression of either H1.1 mutant variant showed a normal occurrence of these meiotic markers, like the control (**Figure 4F-G**). The large companion cells observed in premeiotic ovule primordia persisted at to the tetrad stage (**Figure EV4E**).

These results indicate that impairing H1.1 depletion in SMCs moderately affects heterochromatin reorganization and SMC growth but does not interfere with progression through meiosis.

### Premeiotic expression of the persistent variants H1.1^R57K^ or H1.1^6GC^ differentially impacts gametogenesis

Female meiosis results in the production of four haploid spores, three of which degenerate. The surviving spore enlarges to become the functional megaspore. The functional megaspore will develop into the female gametophyte, or embryo sac, typically through three rounds of endomitosis that are followed by cellularization to form the two female gametes and accessory cells (Skinner & Sundaresan, 2018). Given that expression of H1.1 mutant variants in SMCs does not impact meiosis, we asked whether it affects functional megaspore differentiation and embryo sac development. To this aim we analyzed the functional megaspore-specific marker, *AKV::H2B-YFP,* a fluorescent H2B reporter driven by the *ANTIKEVORKIAN* (*AKV*) promoter (Rotman *et al*, 2005; Schmidt *et al*, 2011). Marker expression showed similar frequencies in wild-type and Dex-treated ovules expressing the mutant H1.1^R57K^ or H1.1^6GC^ variants (**Figure 5A, Figure EV5A**). Embryo sac development, however, was compromised, with a high proportion of embryo sacs at stages FG2-FG4 in ovules expressing either mutant variant whereas in the control, the majority of embryo sacs were mature at stage FG6 (**Figure 5B, Figure EV5B**). To determine whether this was due to developmental arrest or a delay, we emasculated flowers at 9 dpi and allowed them to develop further, re-analyzing them at 11 dpi. Interestingly, in lines expressing the H1.1^R57K^ variant, the proportion of mature embryo sacs at 11 dpi was no longer different than in the control line (**Figure 5B**) suggesting that the abnormal embryo sacs initially detected at 9 dpi were delayed in their development. The same observation was made in lines expressing *amiR[AIH]* with a large fraction of delayed embryo sacs at 9 dpi that reached maturity at 11 dpi (**Figure EV5C**). In contrast, a large proportion of abnormal embryo sacs persisted in lines expressing the H1.1^6xGC^ variant (**Figure 5B, Figure EV5B**).

**Figure 5.**
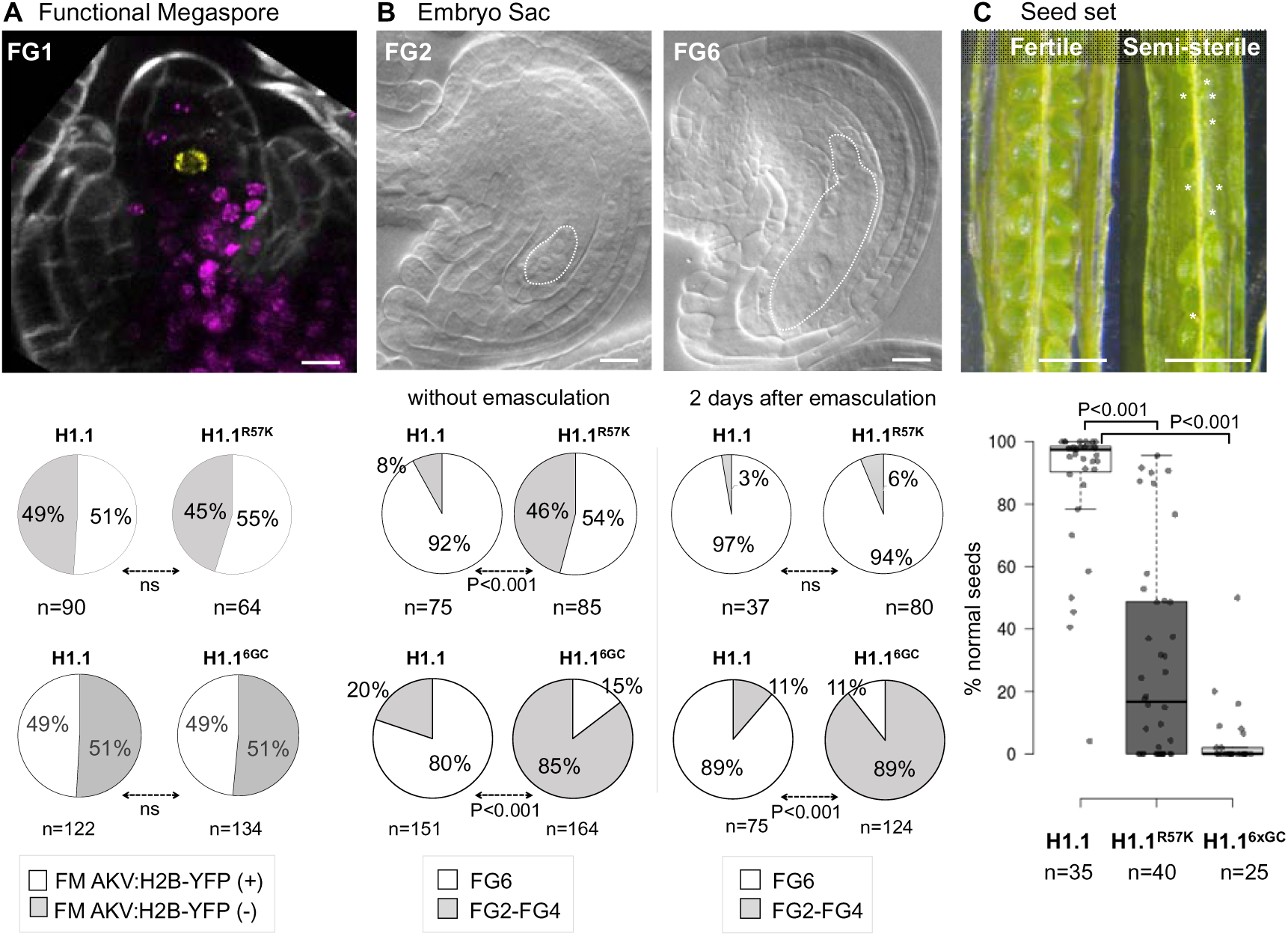
H1.1 persistence in the SMC impairs gametogenesis. **(A-B)** Analysis of functional megaspore (FM) fate establishment and embryo sac development in ovules derived from primordia induced for the expression of the H1.1^R57K^-RFP and H1.1^6xGC^-RFP persistent variants or the H1.1-RFP control variant. Representative images of an ovule at the FG1 stage counterstained with Renaissance (grey) expressing the functional megaspore-specific marker, *AKV::H2B-YFP* (Schmidt *et al*, 2011)(yellow) and H1.1-RFP (magenta) at 7 dpi (A) and of cleared ovules at the FG2 and FG6 stage showing a 2-nucleate and a mature embryo sac, respectively at 9 dpi (B) with respective scoring in F1 plants showing segregation of *AKV::H2B-YFP* (pie charts below) of the functional megaspore marker occurrence at 7 dpi (A) or of mature embryo sacs (FG6) vs earlier stages (FG2-FG4) in ovules at 9 dpi without emasculation or at 11 dpi and after emasculation as indicated. Scoring for each mutant variant-expressing lines was compared to the control line in independent experiments. Slight differences in ovule development progression at the time of flower bud collection between experiments explain the different frequencies of AKV::H2B-YFP expression between replicates scoring in the control line. n, number of ovules scored. P values from a Fisher exact test. Scale bar, 20 μm. **(C)** Seed set analysis in siliques 20 dpi derived from flower buds stage 9 induced for H1.1-RFP, H1.1^R57K^-RFP and H1.1^6xGC^-RFP expression. Representative images of a fully fertile silique (H1.1-RFP control line), and semi-sterile silique (H1.1^R57K^-RFP mutant line) at 20 dpi and scoring of normal seeds (plots) for all three genotypes as indicated. *, infertile ovules. n, number of siliques scored (ca 50 seeds p. silique). P values, Mann Whitney U test. Scale bar: 1mm. See replicate measurements Figure EV5 and raw data in Data Source 5.

The different outcomes we observed in the above experiments suggest distinct functional impacts of the R57K and K89R substitutions in H1.1 on embryo sac development.

### Premeiotic expression of either H1.1^R57K^ or H1.1^6GC^ variant induces seed sterility

We then asked whether the delayed embryo sacs formed following the induction of the H1.1^R57K^ variant in pre-meiotic flower buds were functional. Too this aim, we assessed fertility by scoring the number of plump, viable versus aborted seeds in mature siliques at 20 dpi. The control line induced for the expression of H1.1-RFP produced 87% viable seeds (n=35 siliques, **Figure 5C**). The line expressing H1.1^R57K^-RFP, by contrast, exhibited a highly variable rate of seed abortion ranging from 0% to 95% with an average of 29% viable seeds (n=40 siliques, **Figure 5C**). To explain this high level of seed abortion despite the production of delayed, but morphologically normal embryo sacs, we measured pollen viability. We observed 12% aborted pollen (n=323) in anthers expressing H1.1^R57K^-RFP at 9dpi compared to 3% in the control line (n=425, **Figure EV5D**). This pollen abortion rate cannot explain the high frequency of aborted seeds. Hence, whether the induction of the H1.1^R57K^ variant affects gametic functionality or that of sporophytic tissues remains to be investigated. In contrast, plants expressing H1.1^6xGC^-RFP in premeiotic flower buds, which produce close to 90% abnormal embryo sacs had only 4.5% viable seeds on average (n=40 siliques, **Figure 5C**) and a 67% aborted pollen (n=425, **Figure EV5D**). Thus, in the case of the H1.1^6xGC^ variant sterility is largely due to female and male gametophytic defects although a minor contribution of sporophytic effects cannot be excluded.

Furthermore, the induction of *amiR[AIH]* and *amiR[Cul4]* resulted in high seed sterility compared to mock controls (**Figure EV5E**). This is likely because the AIH and CUL4 enzymes regulate multiple targets relevant to reproduction and not just H1. This is consistent with the embryonic lethality of the *aih* loss-of-function mutant (*emb1873*, Meinke, 2020) and ovule and seed development defects in the *cul4* loss-of-function mutant (Dumbliauskas *et al*, 2011).

## DISCUSSION

The depletion of canonical H1 linker histones is a hallmark of the somatic-to-reproductive fate transition in both plants and animals (She *et al*, 2013; Hajkova *et al*, 2008). In mice, H1.2 depletion in PGCs is critical to establish pluripotency (Christophorou *et al*, 2014). In the flowering plant *Arabidopsis*, the loss of H1.1 and H1.2 in male and female SMCs precedes extensive chromatin reorganization involving both structural alterations and epigenetic modifications (She *et al*, 2013; She & Baroux, 2015) but neither the mechanisms nor the functional implication of H1’s removal were known.

### A citrullination-ubiquitination model for premeiotic H1.1 depletion in female SMCs

Previously, we proposed that the loss of H1.1 in the SMC occurs through proteasome-mediated protein degradation (She *et al*, 2013). Plants possess several pathways for protein degradation but ubiquitin-mediated targeting of proteins to the proteasome is prevalent (Vierstra, 1993; Moon *et al*, 2004). Ubiquitination involves an enzymatic cascade where ubiquitin is transferred from the E1 ubiquitin-activating enzyme to the E2 ubiquitin-conjugating enzyme, and ultimately to the target protein, either directly or indirectly through an E3 ubiquitin ligase (Mazzucotelli *et al*, 2006). In this study, we demonstrate a role for CUL4 in H1.1 degradation in the SMC, extending its previously reported functions in DNA replication, repair, and chromatin remodeling (Biedermann & Hellmann, 2011). Whether CUL4 acts directly or indirectly on H1.1 for instance via the formation of a CUL4 RING ligase (CRL4) complex remains to be determined. CRL4s indeed target nuclear proteins and influence chromatin and chromosome function (Fonseca & Rubio, 2019). For instance, in rice, a CRL4 complex ubiquitinates the Argonaute protein MEIOSIS ARRESTED AT LEPTOTENE1 (MEL1) whose degradation in SMCs is essential for meiosis and the formation of viable microspores (Lian *et al*, 2021).

Presently, profiling histone ubiquitination in Arabidopsis female SMC is technically not possible as of today. But, a previous study identified six ubiquitinated lysines in H1.1 in Arabidopsis seedling tissues (Xue *et al*, 2022), five in the C-terminal tail and one (K89) in the globular domain. Arginine substitutions of these residues as in the H1.1^6xGC^ variant prevented H1.1 degradation in the SMC, with K89 being sufficient to confer this property. These substitutions did not increase H1.1 binding affinity to chromatin, further suggesting a role for PTMs in the degradation process. We propose a model where CUL4 mediates, either directly or indirectly, the ubiquitination of K89 (and possibly other residues) for targeting H1.1 to the proteasome.

Intriguingly, a mutated H1.1 variant (H1.1^11xG^, Figure 1) with 11 substituted lysines residues in the globular domain including the K89R mutation, exhibited normal depletion in the SMC in twelve independent transformants. Sequencing confirmed the presence of all substitutions in the introduced construct (data not shown). While this suggests that K89’s role in degradation requires an intact globular domain, it is unlikely that the 11 K-to-R substitutions disrupt its folding, as this domain is highly resilient to amino acid changes (Martinsen *et al*, 2022). At present we cannot fully explain this observation but suggest that H1.1 degradation may be subject to versatile and redundant mechanisms involving concurrent or cooperative PTMs. In support of this hypothesis, H1.1 in leaves was found to carry a wide variety of PTMs, including crotonylation at six lysines (Kotliński *et al*, 2016).

Furthermore, parallel efforts to identify key amino acids involved in H1.1 degradation pinpointed R57, whose charge matters. Indeed, substituting R57 with a lysine (R57K), which preserves the charge at this position, creates a variant with a similar residency time on chromatin as the wild-type variant. In contrast, neutralizing the charge by an alanine substitution (R57A) creates an H1.1 variant with a higher mobility, i.e a shorter residency time on the chromatin as the wild-type H1.1. Yet, the R57K substitution protects H1.1 from degradation in a large fraction of SMC, the R57A substitution does not. In contrast, the latter seems to induce a faster depletion as suggested by the reduced fraction of SMC with partial depletion at the time of scoring. This finding indicates that H1.1 binding properties play a role in its degradation. In our working model (**Figure 6**), more mobile H1.1 variants, which dissociate faster from chromatin than wild-type variants, for instance due to R57 charge neutralization are more readily targeted for ubiquitination-mediated degradation. Thus, the H1.1^R57A^ variant will be more efficiently degraded than proteins with a slower exchange rate, such as the H1.1^R57K^ variant. The latter degraded at a slower pace should lead to a significant fraction of SMCs with persistent H1.1 (H1.1^R57K^-RFP) signal but will eventually be degraded over time. In line with this model involving first an increased dissociation rate by charge neutralization then degradation, the highly mobile, readily degraded H1.1^R57A^ variant becomes protected from degradation when combined with a K89R substitution (**Figure EV6**).

**Figure 6.**
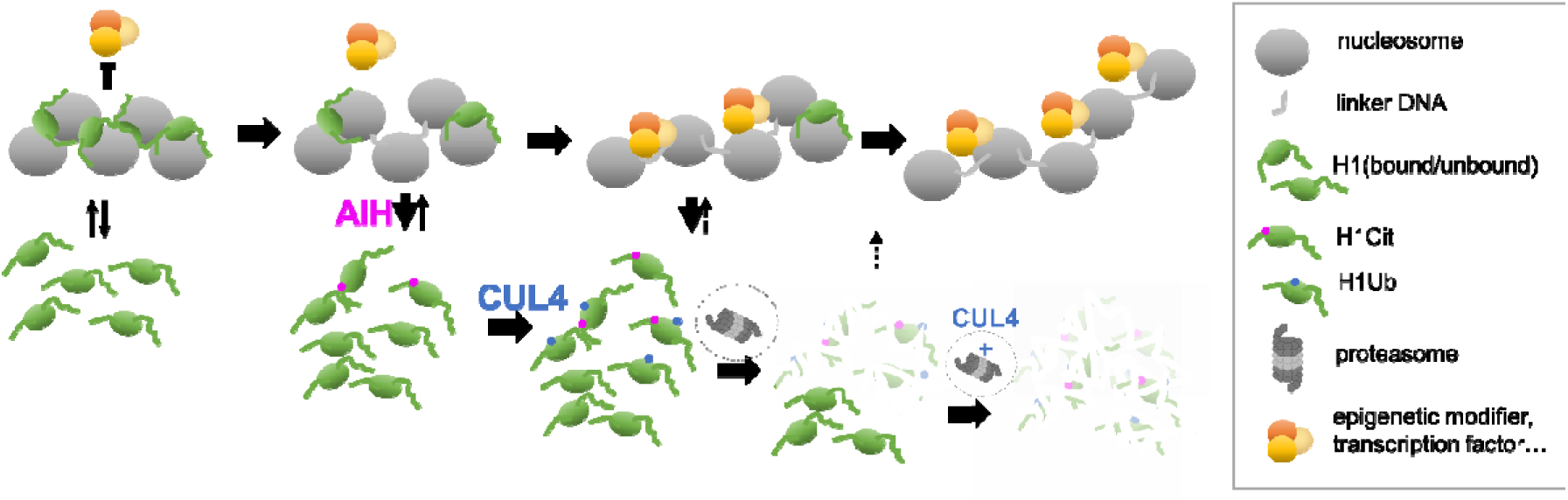
A two-step dissociation-degradation model of H1.1 depletion in the SMC. Based on our findings, we propose a two-step model explaining the depletion of H1.1 from the SMC chromatin. First, the association of H1.1 with chromatin is electrostatically destabilized due to charge neutralization by the citrullinase AIH, particularly at residue R57, while other target residues are not excluded. This facilitates dissociation from chromatin and increases the pool of unbound H1.1 proteins susceptible to ubiquitination by CUL4 and subsequent proteasome-mediated degradation. As H1.1 binding dynamics to the nucleosomes are altered, the progressive degradation of H1.1 depletes the available pool for binding. Meanwhile, reduced transcription and/or translation in the SMC, broad or gene specific, prevents the replenishment of the H1.1 pool. As a result, chromatin becomes accessible to epigenetic modifiers, transcription factors, or both, to render the cell competent for the post-meiotic formation of the embryo sac.

Our model invokes a mechanism that neutralizes the charge at R57. Citrullination, i.e. the deimination of arginine producing a citrulline, results in the loss of a positive charge. In mouse PGCs, this modification is mediated by the citrullinase PADI4, reduces the binding affinity of H1.2 to the nucleosome and was proposed to promote H1.2 depletion in PGCs. Our findings that the plant citrullinase AIH is critical for H1.1 depletion in the SMC support the idea of a similar mechanism in plants (**Figure 6**)

Whether additional residues are modified by AIH, or other enzymes affecting H1.1 mobility, remains to be determined. For instance, the arginine residue R79 located 12 amino acids downstream R57 is conserved in mouse and *Arabidopsis* H1 variants (**Figure EV2D**) and hence another candidate. Additionally, phosphorylation is known to be a major destabilizing PTM of H1 at the onset of S-phase, and is crucial for replication in animal cells (Alexandrow & Hamlin, 2005). The *Arabidopsis* H1.1 variant contains three S/TPxK motifs, which are prone to phosphorylation in the C-terminal tail (Kotliński *et al*, 2016), and CDC2/CDKA;1 can physically interact with H1.1 (Pusch, 2012). It remains to be determined whether this mechanism contributes to H1.1 depletion in the SMC prior to meiotic S-phase (She *et al*, 2013).

The role of citrullination in chromatin regulation has been extensively studied in animal systems, in disease and development. Particularly, histone hypercitrullination is responsible for chromatin decondensation in activated neutrophils upon inflammatory immune response and for the production of chromatin-containing neutrophil extracellular traps (Maronek & Gardlik, 2022; Zhu *et al*, 2023). PAD enzymes are also associated with oncogenicity where their ectopic activation leads to chromatin decondensation, ectopic histone citrullination and deregulation of gene expression (Zhu *et al*, 2021; Zheng *et al*, 2020). Yet, histone citrullination also plays a role in development. Notably, the citrullination of H1 (Christophorou *et al*, 2014) but also of H3 and H4 (Xiao *et al*, 2017) are necessary for activating pluripotency genes in PGCs. After fertilization, selective citrullination of H3 and H4 is necessary for embryonic genome activation and proper gene expression (Zhang *et al*, 2016).

By contrast, much less is known about citrullination in plants. The catalytic domain of the plant AIH citrullinase is highly conserved with its mammalian PAD4 counterpart: it shares the four key catalytic amino acids and exhibits similar inhibitor sensitivity, indicating that citrullination may play a fundamental role in plants too. Indeed, protein citrullination has recently been identified as a significant post-translational modification in response to cold stress in cell cultures (Marondedze *et al*, 2021). Our discovery that AIH is required for H1.1 depletion in SMCs opens new avenues for investigating the role of histone citrullination in plant chromatin dynamics.

A key question arising from our findings is the cell specificity of H1.1 degradation. Both AIH and CUL4 are broadly expressed in ovule primordia, a pattern that does not explain the observed cell-specific degradation. One possibility is that these enzymes might be catalytically regulated by a cofactor conferring cell specificity. For example, mammalian PADI4 activity is influenced by Ca²[ availability (Arita *et al*, 2004). In an alternative model, increased H1.1 dissociation followed by degradation may be the default process in all cells of the primordium but low or stalled transcription and/or translation in the SMC could prevent replenishment of the H1.1 pool. With an average polyA tail of ∼55-60 nt and a half-life of about 1 hour in vegetative tissues, H1.1 transcripts do not exhibit distinctive features (Jia *et al*, 2022; Szabo *et al*, 2020). However, the short lifetime of H1.1 transcripts in the SMC progenitor, in combination with a relative transcriptional quiescence (She *et al*, 2013) may be sufficient to prevent the replenishment of H1.1 on chromatin. Whether translational inhibition also occurs in the SMC, targeting specific transcripts such as those encoding H1.1, or if it is part of a broader mechanism —similar to what is seen in animal PGCs (Oulhen *et al*, 2017)— remains to be determined. Interestingly, premeiotic spikelet and male meiocytes in prophase I of rice are enriched in phased secondary small interfering RNAs (phasiRNAs) involving MEL1 (Komiya *et al*, 2014) and bearing the potential for translational inhibition (Jiang *et al*, 2020; Liu *et al*, 2020). Future studies could explore the repertoire of small and antisense RNAs in the female SMCs of *Arabidopsis*. This model, in which H1.1 undergoes rapid turnover throughout the primordium but is not replenished in the SMC due to stalled transcription and/or translation may also apply to other chromatin constituents such as H2A.Z and H3.1/HTR13 that are specifically lost in the SMC (She *et al*, 2013; Hernandez-Lagana & Autran, 2020). This opens the exciting question of which process(es) specifically reduce transcriptional and/or translational activity in the SMC.

### Implications of premeiotic H1.1 clearance for reproduction

H1 depletion in mouse PGCs and subsequent chromatin reorganization are essential for establishing pluripotency in the mammalian germline (Christophorou *et al*, 2014). Although similar events were known to occur during SMC formation in *Arabidopsis*, marking the transition from somatic to reproductive fate (She *et al*, 2013; She & Baroux, 2014), the precise role of H1 depletion remained unknown. Since linker histones interplay with the deposition and maintenance of various epigenetic marks in *Arabidopsis*, including DNA methylation, histone methylation (including H3K27me3), histone acetylation, and intergenic antisense transcription (Wierzbicki & Jerzmanowski, 2005; Rutowicz *et al*, 2019; Zemach *et al*, 2013; Choi *et al*, 2020), their depletion at the onset of SMC differentiation likely opens a window of opportunity for epigenetic reprogramming. With our current knowledge derived from H1 loss of function mutants, we had expected that SMCs compromised in H1.1 depletion e.g., due to expression of either the H1.1^R57K^ or H1.1^K89R^ variants, would show incomplete chromatin reorganization. In this scenario, we expected maintenance of heterochromatin instead of the decondensation normally seen in SMCs (She *et al*, 2013), and similar levels of CHH methylation and H3K27me3 in SMCs and surrounding cells, rather than the decrease observed in SMCs (She *et al*, 2013; Ingouff *et al*, 2017). We detected mild differences in the H1.1^R57K^-RFP expressing line, and little to no effect when expressing H1.1^K89R/6xGC^-RFP. These distinct outcomes can be explained by our two-step dissociation-degradation model (**Figure 6**). In this model, the unbound pool of H1.1^R57K^ variants form at a slower rate than citrullinated H1.1^K89R/6xGC^ variants and, by contrast to the latter that are protected from degradation, H1.1^R57K^ variants are degraded but at a slower rate than the wild-type H1.1 variant.

We found that H1.1 depletion in SMCs is not essential for meiosis but is crucial for establishing the developmental competence of the functional megaspore to form a functional embryo sac. We propose that premeiotic H1 depletion plays a critical role in establishing an epigenetic landscape in the SMC that prepares the post-meiotic transcriptional program in the functional megaspore, thus controlling its developmental competence. Since the functional megaspore generates a multicellular embryo sac comprising different cell types, it is pluripotent. Thus, similar to H1 depletion in mouse PGCs, H1 depletion in Arabidopsis SMCs seems necessary to establish pluripotency in the functional megaspore. However, this hypothesis requires further investigation, e.g., through transcriptome analyses of functional megaspores expressing the different H1.1 variants.

Functional megaspores derived from SMCs where H1.1 depletion was disrupted, were compromised in mitosis, resulting in delayed or arrested embryo sac development at the 2-4 nucleate stage. This finding suggests a failure in the expression of one or several components controlling endomitosis, nuclear migration in the syncytium, cell growth, or a combination thereof. Notably, the H1.1^R57K^ mutant, affecting H1.1 dissociation rate from chromatin in the SMC, had a different impact on embryo sac development than the H1.1^6xGC^ mutant, which abolishes degradation. In the former, embryo sac development was delayed, while in the latter, it was arrested. These findings could be explained by a scenario where although H1.1^R57K^ persists in the SMC, a relevant fraction of H1.1 still undergoes degradation, enabling some degree of epigenomic reprogramming. In contrast, the continuous presence of the H1.1^6xGC^ mutant, which cannot be degraded, compromises epigenomic reprogramming more severely.

Overall, our study sheds light on the essential role of H1 regulation prior to meiosis to establish post-meiotic competence that is necessary for embryo sac developmental and thus reproductive success. The parallels observed between plant and animal systems in the regulation of histone H1 dynamics highlight a potential evolutionary convergence in the mechanisms governing reproductive cell fate determination. In both plants and animals, the removal of histone H1 and other chromatin modifications is crucial to establish a chromatin landscape conducive to cell fate transitions (Hajkova *et al*, 2008; She *et al*, 2013). Understanding the underlying mechanisms in plants can inform strategies to manipulate reproductive processes in crops, potentially improving fertility and yield.

## METHODS

### Plant growth conditions

*Arabidopsis* seeds were sterilized in freshly made 0.03% bleach and 0.05% Triton X-100 for 10 min, washed three times in sterile water, briefly incubated in 70% EtOH and washed once before sowing on the germination medium consisting of 0.5xMurashige and Skoog salts (MS, Carolina 19-5700, USA), 10% (w/v) Bactoagar, pH 5.6. Seeds were stratified 2 to 4 days at 4°C before being transferred to a growth incubator (Percival) with long day conditions (16h light [120μE m-2s-1] at 21°C and 8h dark at 16°C). Ten days-post-germination, seedlings at the 2-4 leaves stage were transplanted to soil, covered with a layer of sand and imbibed with Solbac (Andermatt Biocontrol, Switzerland) for pest biocontrol. Plants were cultivated in a growth chamber with controlled conditions under a long-day photoperiod (16 hours light) at 18-20°C and 38% to 59% rF of humidity.

### Generating construct and transgenic lines

The artificial micro-RNAs (amiRNA) targeting CUL4 (At5g46210) or AIH (At5g08170) were designed with the wmd3 database (Meyers & Green, 2010) and comprised the following sequences: 5’-ATGC*TCATGATTCGTAGTATA-3’ and 5’-ACGC*GAGCCGTTCATTAATTA*-3’, respectively, with mismatches to the original sequence indicated by an *. The amiRNAs and the H1.1 mutant variants modified for specific amino acid substitutions were synthesized by Genscript (GenScript Biotech Corporation, genscript.com) and introduced into the *BAR* gene-containing donor vector pRPS5a::LhGR2-GUS::pOP6::attL1-cddb-attL2 vector (Samalova *et al*, 2019) via Gateway cloning (Thermo Fischer Scientific, USA) according to the manufacturer’s recommendation. The resulting constructs, abbreviated as *amiRNA[Cul4]* and *amiRNA[AIH]*, were transformed via *Agrobacterium tumefaciens* (GV3101) into a *pH1.1::H1.1-GFP; pH1.2::H1.2-ECFP*; *3h1 Arabidopsis* line. This line was created by introgression of *pH1.1::H1.1-GFP* and *pH1.2::H1.2-ECFP* into the triple H1 mutant described (She *et al*, 2013) by crossing and selection. The *pH1.1::H1.1-GFP* line was described (Rutowicz *et al*, 2015) and the *pH1.2::H1.2-ECFP* line was created as described for the *pH1.2::H1.2-GFP* line (Rutowicz *et al*, 2015) albeit replacing the *GFP* sequence with the *ECFP* sequence (Clontech).

The constructs encoding RFP-tagged H1.1 variants containing codon changes for amino acid substitutions were synthesized by Genscript (GenScript Biotech Corporation, genscript.com), introduced into the same donor vector via Gateway cloning as described above and transformed into the *Arabidopsis* Col-0 accession. Positive T1 plants were identified based on selection for the BAR gene *in vitro* (0.5x MS medium supplemented with 50 μM phosphinothricin) followed by a GUS reporter assay on the 3^rd^ or 4^th^ true leaf after treatment with 10 μM dexamethasone (see below).

### Histochemical detection of GUS reporter activity (GUS reporter assay)

To select transgenic lines with a functional construct we verified each resistant T1 plants for the expression of the *uidA* reporter gene encoding a β-glucuronidase (GUS) and present in the responder cassette (Craft *et al*, 2005; Samalova *et al*, 2019). For this, one cauline leaf per seedling was incubated with the reaction buffer (0.1% Triton X-100, 10mM EDTA, 2mM ferrocyanide, 2mM ferricyanide, 100mM Na_2_HPO_4_, 100mM NaH2PO4, 2mg/mL X-Gluc) in 48-multiwell plates, vacuum infiltrated for 5min and stained for 6-8 hours at 37°C.

### Dexamethasone treatments

*Arabidopsis* lines containing the dexamethasone (Dex)-inducible constructs were induced as described (Schubert *et al*, 2022). Briefly, flower buds were delicately opened with dissecting needles under the stereomicroscope to allow for a good exposure of the carpel, then gently brushed, using a fine paint brush with soft hairs, either with the induction solution (10 μM Dexamethasone [Sigma-Aldrich Merck KGaA, Germany], 0.1 % DMSO [Sigma-Aldrich Merck KGaA, Germany], 0.01% Silwet-L77, ddH2O) or with a mock solution (0.1 % DMSO, 0.01% Silwet-L77, ddH2O). Flower buds or siliques were collected several days post-induction (dpi) depending on the experiment. The preparation of this material for microscopical analyses is described further below.

### Epoxomycin and Cl-amidine treatments

For epoxomicin treatment, intact inflorescences were collected and flower buds older than stage 10 (Smyth *et al*, 1990; Yu *et al*, 2020) were removed. The remaining inflorescences were immersed in an epoxomicin-containing solution (5μM epoxomicin (LUCERNA-CHEMÒ), 0.01%Silwet-77L, 0.01%Tween-20, ddH_2_0) in a 48 multiwell-plate and incubated in a growth incubator (Percival, long days, 21°C) for 2 days before sample preparation for imaging. For Cl-amidine treatment, young flower buds were treated with 1 μM Cl-amidine (CalbiochemÒ) supplemented with 0.01% Silwet-L77 or with a mock solution (0.01 %DMSO, 0.01% Silwet-L77) the same way as for the Dex treatments described above.

### Western-blot and quantification

For protein extraction, ∼100mg of 10 days old *Arabidopsis* seedlings were harvested and ground using a Retsch homogenizer in liquid nitrogen. The ground tissue was resuspended in 400 µL of SDS lysis buffer (40 mM Tris-HCl pH 7.5, 10% glycerol, 5 mM MgCl2, 4% SDS). The samples were incubated at room temperature for 10 minutes under gentle shaking at 400 rpm (Eppendorf ThermoMixer) to solubilize the tissue evenly in the SDS lysis buffer. Following a centrifugation at 13’000g for 5 min the supernatant was collected, and centrifugated again at 13’000g for 5 min. The final supernatant was collected for protein quantification using the Pierce BCA Protein Assay kit (Thermo Fisher Scientific), following the manufacturer’s protocol with 10 µL of the sample/standard. All sample concentrations were normalized by diluting with SDS lysis buffer. The freshly prepared protein samples were then mixed with 1 µL 5x loading buffer (250 mM Tris-HCl pH 6.8, 5% SDS, 50% glycerol, 1 mg bromophenol blue, 5% β-mercaptoethanol) and boiled for 10 min at 95°C.

For immunoblotting, 10 µg of protein extract was loaded for each sample onto an SDS page gel (Mini-PROTEAN TGX™ Precast Gels, Bio-Rad, #4561094). Electrophoresis was performed at 90V, followed by the transfer of proteins onto a nitrocellulose membrane using the Trans-Blot Turbo Transfer System (Bio-Rad) at 2.5A constant, 25V for 20 min. The membrane was stained 1min with Ponceau red (PanReac AppliChem, Cat#A2935,0500), before acquiring an image with the ChemiDoc Imaging System (Bio-Rad). The membrane was then washed with ddH_2_O and blocked with TBS-T (1xTBS, 1% Tween) blocking buffer containing 5% bovine serum albumin (BSA) with shaking for at least 60 min. The membrane was sealed in a plastic pouch and incubated overnight at 4°C on a shaker with 3 mL of primary anti-CUL4 antibody (PhytoAB Cat#PHY0850S) at 1:2000, all diluted in blocking buffer.

Following the primary antibody incubation, the membrane was washed three times for 10 min each with TBS-T before incubation with the secondary HRP-conjugated anti-rabbit IgG antibody (Amersham NA934V Cat#17550629) diluted at 1:10,000 in TBS-T supplemented with 5% milk, for 1h on a shaker. The membrane was washed again three times for 10 min each with TBS-T then incubated for 1 min with 500 µL of Clarity Max Western ECL Substrate (Bio-Rad 1705062) and imaged using the ChemiDoc Imaging System (Bio-Rad). The relative intensity mean was calculated for each sample as the ratio between the intensity mean of the predicted band around CUL4 ∼91KDa and the intensity mean of the major band on the Ponceau staining ∼70kDa. The ratio was then expressed relative to Line4-Rep2 set to “1”. Measurements were done using Fiji on 8bit greyscale inverted images using ROI of the same size.

### 3D Confocal imaging of whole-mount ovule primordia

Carpels freshly dissected from flower buds were delicately opened using insulin needles to remove the carpel walls and expose the ovule primordia, and mounted in Renaissance solution (final concentrations: 1:2000 Renaissance;10% glycerol; 0.05% DMSO; 0.1%Triton X-100 in 1x PBS (modified from Musielak *et al*, 2015) on a Superfrost^TM^ slide (Thermo Fisher Scientific) and covered with a 18×18mm coverslip, #1.5 thickness (Assistant, Germany) and immediately imaged. Fixed, cleared ovules stained with propidium iodide in acrylamide pads (see below) were mounted in Vectashield (H1300-10, Vector Laboratory, USA) and covered with a 18×18mm coverslip, #1.5 Thickness (Assistant, Germany). Serial image acquisition was carried out using confocal microscopy (CSLM Leica SP5 Resonance, or Stellaris, Leica Microsystems, Germany) with an HC PL APO 63x/1,30 GLYC CORR CS2 objective lens, using the fluorophore’s optimal excitation laser, an emission window of circa 40-60nm centered on the emission maximum, using a resonance scanning mode (8000Hz), photon counting mode for detection, and background free HyD detectors, with signal accumulation per line to optimize the pixel intensity distribution. Images were acquired with voxel size of 0.08-0.1μm x,y,z dimension.

### Microscopy-based scoring of the H1 depletion phenotype in female SMCs

The H1.1 depletion phenotype in SMCs was scored on several replicate ovule primordia from independent flower buds under live scanning (see above). Three categories were considered: (i) Full depletion: no detectable signal in the SMC; (ii) Partial depletion: detectable signal, usually in chromocenters; (iii) Persistence: clear signal in both euchromatin and chromocenters.

### Microscopy-based scoring of ovule stage-specific markers

Ovule primordia expressing the SMC, meiosis and gametophyte specific markers *KNU::nlsYFP* (Tucker *et al*, 2003), *ASY1::ASY1-YFP*(Valuchova *et al*, 2020) and *AKV::H2B-YFP* (Rotman *et al*, 2005) respectively, were prepared as described for microscopy analysis. Ovules were scored under live imaging for presence/absence of signal at developmental stages and in genetic backgrounds indicated in the Figures and associated data.

For tetrad analysis in ovules, the samples were mounted in Renaissance solution (final concentrations: 1:2000 Renaissance;10% glycerol; 0.05% DMSO; 0.1%Triton X-100 in 1x PBS (modified after Musielak *et al*, 2015). The presence of normal versus abnormal tetrads was scored under live, 3D imaging at stages and in genetic backgrounds indicated in the Figures and associated data.

### Analysis of embryo sac development

Embryo sac development was analyzed by identifying the developmental stages as described (Schneitz *et al*, 1995), either using the *AKV::H2B-YFP* reporter line (Schmidt *et al*, 2011) introgressed into the mutant backgrounds, or by tissue clearing as indicated in the relevant Figures. For tissue clearing, inflorescences were fixed in acetic acid: EtOH solution (3:1) and incubated for at least 4 hours at room temperature. The samples were then transferred in 70% EtOH and mounted in a clearing solution (chloral hydrate: water: glycerol 8:2:1 w/v/w). The samples were analyzed using transmission light microscopy (Leica DMR, Leica Microsystems, Germany) with differential interference contrast (DIC), using a 20x or 40x dry objective (NA 0.75 and 0.5). Images were acquired using a digital camera (Flexacam C3 Leica Microsystems, Germany).

### Whole mount propidium iodide (PI) staining

Whole-mount staining of ovule primordia was done as described (She & Baroux, 2014). Briefly, Dex-treated flower buds collected at 5 dpi were fixed in 1% formaldehyde,10% DMSO in PBS-Tween (0.1%), then dissected and embedded in 5% acrylamide pads on microscope slides. Tissue processing included clarification, cell wall digestion and permeabilization before application of 10μg/ml propidium iodide (PI). Samples were mounted in Vectashield supplemented with PI (Invitrogen)

### Image processing for signal quantification, cell volume measurement, and RHF analysis

Image processing for 3D reconstruction, segmentation, signal, or cell size segmentation was performed with the Imaris software (Bitplane, Switzerland). Images presented in the Figures correspond to 3D reconstruction or partial z-projections, using orthogonal or oblique slicers, encompassing the SMCs and surrounding cells.

Cell segmentation for cell size measurements was done as described using ImarisCell (Mendocilla Sato & Baroux, 2017) based on the Renaissance signal in cell walls (Musielak *et al*, 2015). Cell volumes of the SMC, the companion cell, and the neighboring cells were exported for analysis.

For quantifying fluorescent reporter signals in nuclei, the later were segmented using Imaris’ surface tool, in the semi-automated segmentation mode guided by user-defined manual contours. The sum of voxel intensities (intensity sum) per channel and per nucleus was exported for analysis. If not otherwise indicated, the relative intensity in the SMC was expressed as a ratio between the intensity sum in the SMC nucleus divided by the average intensity sum of 3-4 nuclei from neighboring L2 nucellar cells.

The relative heterochromatin fraction (RHF) was calculated as the sum of intensity sums in the chromocenters (CC) divided by the intensity sum in the nucleus. Measurements were done on intensity sum projections encompassing the SMC and surrounding nucellus nuclei, processed in Fiji for manual regions-of-interest (ROI) definition.

### 3D protein structure alignment

The 3D predicted protein structures 1VKP for AIH (green) and 2DEW for PADI4 (yellow) were superimposed in COOT (Emsley *et al*, 2010) and the result was visualized in PyMOL (Chaudhari & Li, 2015).

### Fluorescence Recovery After Photobleaching (FRAP)

Measurements were done on root tips of five days old seedlings essentially as described(Rosa, 2018) using a confocal laser scanning microscope equipped with a FRAP module (Leica SP8-R, inverted, Leica microsystems, Germany). Briefly, one sample was prepared at a time, with one root gently mounted (without squashing) in freshly made liquid 0.5x MS. The slide was sealed with transparent nail polish and let to equilibrate for 5-10 min on the microscope stage, with the imaging chamber set at a constant temperature of 20°C. Bleaching and imaging were done using an APO PL 63x water immersion objective, NA 1.4, over a single plane with the pinhole increased up to maximum (5.38 AU) to capture the entire nucleus. Bleaching was performed within a region-of-interest (ROI) of 2μm diameter, positioned in euchromatin, using 3-5 pulses to reach near full bleach of the signal, using the maximal power and full transmission of the 561nm laser. Post-bleach images were recorded using low laser intensity for sustainable acquisition, with two sequences: the first sequence of 30 images with 265ms intervals captures the initial, rapid recovery phase; the second sequence captured the slow recovery phase with 24 images with 10 sec intervals. For analyzing fluorescence recovery, images were first corrected for nuclear drifts occurring during acquisition, using a rigid registration approach in Fiji (Schindelin *et al*, 2012). Fluorescence measurements were done for the bleached ROI, a background ROI of the same size outside the nucleus, and for the nucleus captured manually with contours. The calculation of fluorescence recovery using double normalization was done as described (Rosa *et al*, 2015) with intensities at each time point expressed relative to the initial intensity (becoming 1, maximum intensity before bleaching) for each image for comparison.

### Fertility analysis

Single inflorescences were induced with either 10μM Dex or a mock solution as previously described. At 19-21 dpi, the 6^th^, 7^th^, 8^th^ silique of each treated plant was collected, and the number of infertile ovules (white, shriveled), aborted seeds (brow, shriveled) and viable seeds (green, plump) was scored.

### RNA *in situ* hybridization

The antisense RNA probe was designed to target all splicing versions of the *AIH* transcripts (At5g08170) and covers position 441-774 of the coding sequence (see **Figure EV3B**). RNA probes were synthesized and labelled using the DIGOXIGENIN (DIG) RNA Labeling Kit (DIG RNA Labeling Mix, Cat.No. 11277073910, Sigma-Aldrich Merck KGaA, Germany) following the manufacturer’s instructions. Probe synthesis was performed using the following forward and reverse primers 5’-CGCTTTGGAACGAATTCC-3’ and 5’-TAATACGACTCACTATAGGGCCGAAAGAGCTTCCACAG-3’, respectively. The *in vitro* transcription reaction was carried out using the T7 RNA polymerase according to the manufacturer’s recommendation (Sigma-Aldrich Merck KGaA, Germany). RNA probe preparation and *in situ* hybridizations were done as described (Dreni *et al*, 2007). Briefly, single inflorescences (Col_0) were fixed in ice-cold fixative (4% paraformaldehyde (w/v), 1xPBS), embedded in wax using a tissue processor (ASP200, Leica BIOSYSTEMS, Switzerland AG). 6-µm-thick sections were prepared on Superfrost^TM^ Plus Adhesion Microscope Slides (Thermo Fisher Scientific Inc), dewaxed using Histoclear (Sigma-Aldrich Merck KGaA, Germany) and dehydrated using a series of EtOH dilutions. Following Proteinase K digestion, slides were fixed in formaldehyde solution, treated with acetic anhydride and dehydrated using a series of EtOH dilutions. RNA hybridization was performed overnight at 43°C in 50% formamide, 2xSSC. Post-hybridization steps were carried out with a series of formamide dilutions, before washes and dehydration using an EtOH dilution series. Detection was carried out with an anti-digoxigenin antibody conjugated with alkaline phosphatase (AP), 1:700 (Anti-Digoxigenin-AP, Fab fragments, Sigma-Aldrich Merck KGaA, Germany). The chromogenic reaction was performed overnight using the BCIP alkaline phosphatase substrate and NBT catalyst (BCIP/NBT Color Development Substrate, Promega, Switzerland). Slides were analyzed under transmission light microscopy (Leica DMR, Leica Microsystems, Germany) and images were acquired with a digital camera (Flexacam C3, Leica Microsystems, Germany).

### Measurement of pollen viability

Flower buds at stages 5-7 were treated with Dex as before and flowers at stage 8– to 9 were collected 9 days post induction (9 dpi). Anthers containing mature pollen were dissected, gently squashed onto slides, and fixed in 10% EtOH for 10 minutes. Samples were then mounted in an Alexander staining solution (10% Ethanol, 0.01% Malachite green, 25% glycerol, 0.05% acid fuchsin, 0.005% Orange G, 4% glacial acetic acid in distilled water (modified after Peterson *et al*, 2010).

## ACKNOWLEDGEMENTS

We thank Diana Pazmino, Valeria Gagliardini, Christoph Eichenberger for technical assistance in plant care, digital qPCR, microscopy, respectively, Sara Simonini and Daniela Guthörl (University of Zurich) for advice with RNA *in situ* hybridization, Stefano Bencivenga (University of Zurich) for advice regarding recalcitrant cloning, Frédérique Pasquer and Ramona Hartman (University of Zurich) for in-house sequencing, Arturo Bolanos and Peter Kopf (University of Zurich) for general lab support, Vimal Rawat (University of Zurich) for help with the expression map of *CULLIN* genes, the central microscopy facility (ZMB) of the University of Zurich for microscopy support, Mathieu Ingouff (IRD Montpellier, France) for the *mCHH-YFP* DynaMet line, Mathew Tucker (University of Adelaide, Australia) for the *KNU::nlsYFP* line, Wei Cai Yang (Institute of Genetics and Developmental Biology, Beijing, China) for the *AKV::H2B-YFP* line, Stefan Heckmann (IPK Gatersleben, Germany) for the *ASY1::ASY1-YFP* line, and Daphné Autran (IRD, Montpellier, France) for critical reading of the manuscript and insightful suggestions.

The project was supported by the University of Zurich, the Swiss National Science Foundation (grants #310030_185186 to CB and #31003A_179553 to UG), and the European Union’s Horizon 2020 research and innovation program under the Marie Sklodowska-Curie (MSC) grant agreement No 847585 (RESPONSE, to CB and SB), and benefitted from networking events of the European Cooperation in Science and Technology (COST) Action # CA16212.

## AUTHOR CONTRIBUTIONS

Conceptualization, C.B.; Methodology, Y.L., D.F., J.S., S.B., U.G. and C.B.; Formal Analysis: Y.L., D.F. and C.B.; Investigation, Y.L., D.F., J.S., K.R., A.L., Z.K. and C.B.; Software: A.G.F.; Writing – Original Draft, Y.L. and C.B.; Writing – Review & Editing, Y.L., D.L., U.G. and C.B.; Visualization: Y.L. and C.B.; Funding Acquisition, S.B., U.G. and C.B.; Resources: U.G., S.B. and C.B.; Supervision, SB, UG and C.B.

