## supplementary figures and legend for "Pre-meiotic H1.1 degradation is essential for Arabidopsis gametogenesis"

Li, Fei, Schubert et al

**Expanded View Figures and legends**

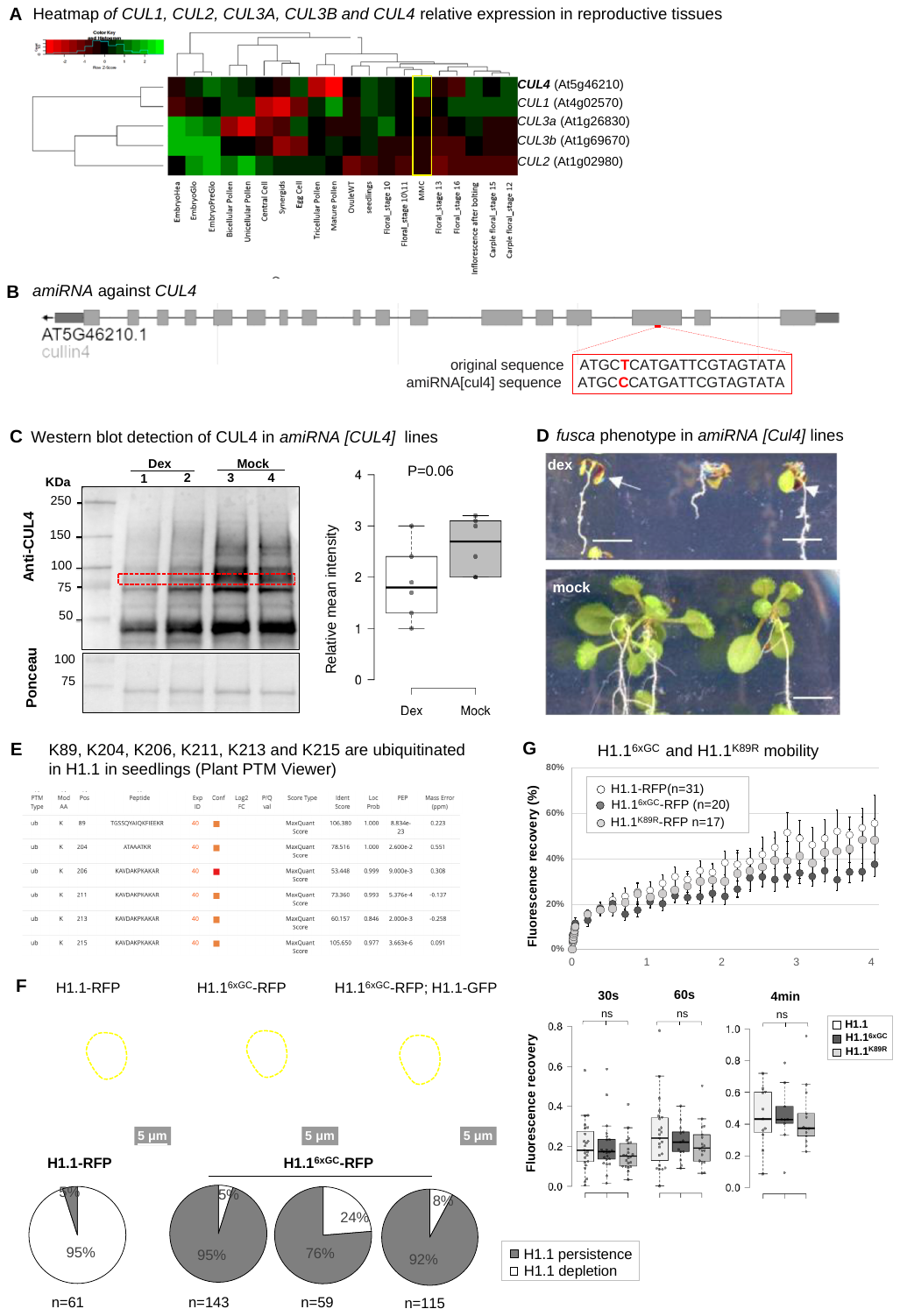

**Figure EV1. H1.1 degradation is controlled by CUL4 and the K89 residue, without altering its stability (related to Figure 1)**

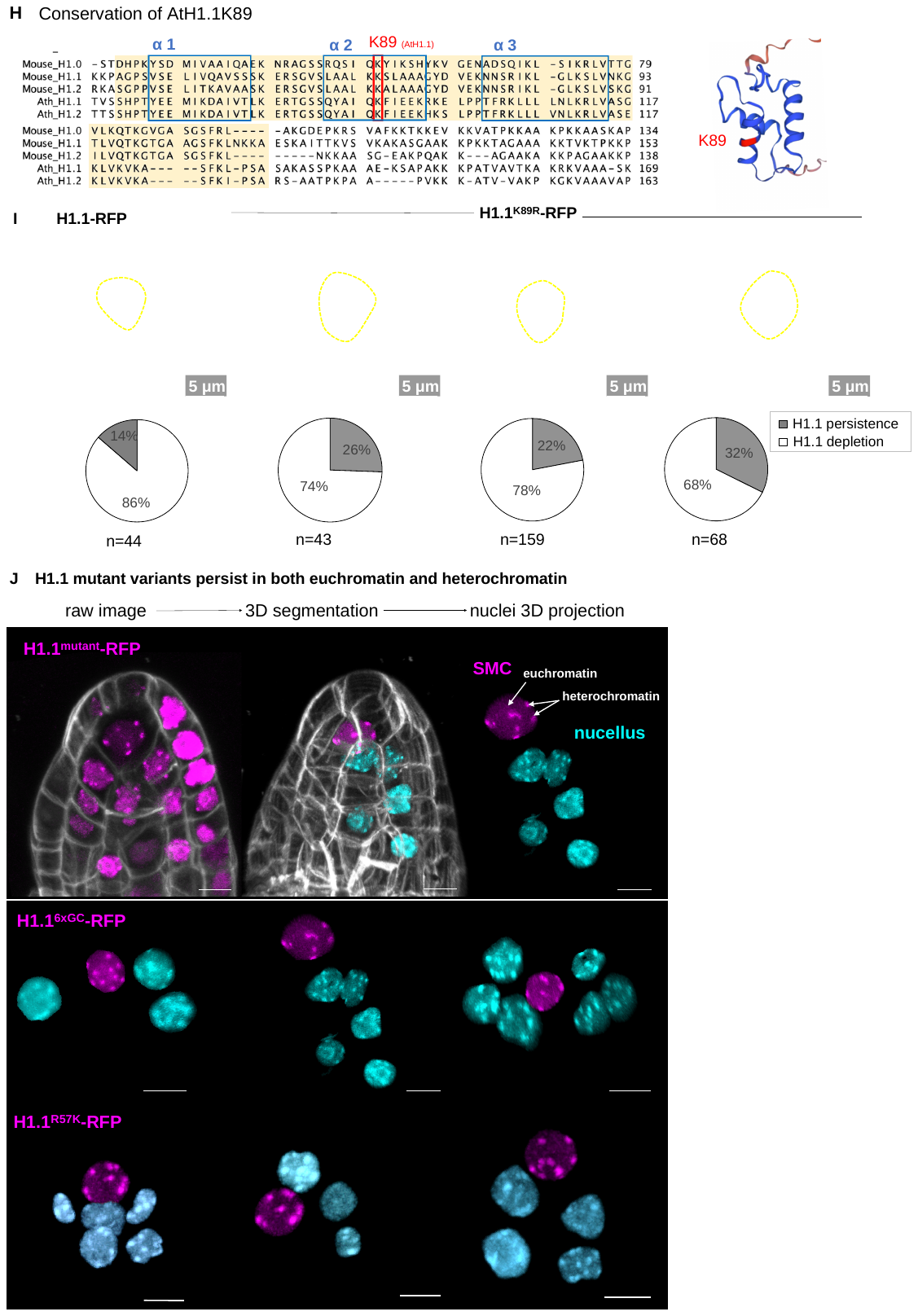

**Figure EV1. H1.1 degradation is controlled by CUL4 and the K89 residue, without altering its stability (continued)**

**Figure EV1. H1.1 degradation is controlled by CUL4 and the K89 residue, without altering its stability (related to Figure 1)**

**(A)** Relative Expression of CUL1 (AT4G02570), CUL2 (AT1G02980), CUL3A (AT1G26830), CUL3B (AT1G69670), CUL4 (AT5G46210) in different tissue present in Affymetrix ATH1 GeneCHIP experiments from publicly available datasets (Wuest *et al*, 2010; Borges *et al*, 2008; Honys & Twell, 2004; Pina *et al*, 2005; Schmidt *et al*, 2011). The dendrogram in the expression heatmap was generated using hierarchical clustering with default options in the heatmap.2 function from the gplots package in R (R Core Team, 2023; Warnes *et al*, 2024). Pairwise distances were calculated using the Euclidean distance metric, and hierarchical clustering was performed using the complete linkage method. These settings were used to cluster both rows and columns. The table with normalized expression values can be found in Dataset EV 1, Tab S4A. Abbreviation: MMC (megaspore mother cell), EmbryoHea (embryo heart stage), EmbryoPreGlo (embryo pre-globular stage). (**B)** Schematic representation of *CULLIN4* (At5g46210) genomic region with indication of the position and sequence of the artificial micro-RNA used in *amiR[CUL4]* lines. (**C**) Western blot (left panel) detection using an anti-CUL4 antibody and Ponceau staining below in *amiRNA [CUL4]* Line #8, treated with Dex and mock, each in two independent replicate extractions. Total proteins were extracted from seedlings at 12DPI, loaded and immunoblotted as described in the Methods. The expected molecular weight of CULLIN 4 is indicated (~91KDa). Right: quantification of the relative CUL4 band intensity (see Methods) in three independent lines (4,#7 and #8) , each in two replicate protein extraction. All the blots used for quantifications can be found in Dataset EV 4, TabS4C (**D**) *CUL4* knockdown as in the Dex inducible *amiRNA[CUL4]* line recapitulates a known f*usca* phenotype (Chen *et al*, 2006). Seedlings are 11DAG-old and were grown on ½ MS supplemented with 10uM DEX (upper panel) or a MOCK solution (bottom panel). Scale bar=5mm. (**E)** Screen shot from the Plant PTM Viewer webserver. The Plant PTM Viewer shows six putative ubiquitination sites on Arabidopsis H1.1 (Willems *et al*, 2019). (**F)** Representative images of H1.1-RFP, H1.1^6xGC^-RFP in a wild-type background and co-expressed with H1.1-GFP under its native promoter (She *et al*, 2013) showing persistence of H1.1^6xGC^-RFP and depletion of H1.1-GFP in the SMC (dashed lines) of ovule primordia 5dpi; Pie charts show replicate measurements of persistence vs depletion categories in independent lines. P value: Fisher exact test. (**G)** Fluorescent Recovery After Photobleaching experiments measuring the mobility of the different H1.1 variants as indicated in seedling roots and Boxplot showing the recovery rate at 30s, 60s- and 4-min. P values, Mann-Whitney U test. (**H)** Alignment of selected Arabidopsis and mouse H1 variants (AtH1.1: AT1G06760 H1.1; AtH1.2: AT2G30620 H1.2, Mouse H1.0: P10922, Mouse H1.1: P43275 , Mouse H1.2: P15864) cropped around the globular domain (yellow) showing the position of the conserved Lysin residue (K89 in AtH1.1) and the three α helices as indicated. Right: representation of AtH1.1 3D folding of the globular domain, indicating the position of K89 (**I)** Replicate measurements of H1.1^K89R^ persistence vs depletion in the SMC of independent lines induced as described in the main text and compared to a control line. P value: Fisher exact test. (**J)** 3D projection showing H1.1^6GC^-RFP and H1.1^R57K^-RFP persistence in both euchromatin and heterochromatin. Top panel: representation of the image processing used to isolate nuclei in silico for projections (left, whole primordium counterstained with Renaissance; middle, primordium after segmentation and masking of the SMC nucleus (magenta) and several nucellus nuclei (cyan); right: nuclei projection only) Middle and bottom panels: three representative images for the H1.1 variants as indicated. Scale bar: 5µm. Original data used in graphs in **Dataset EV 1**

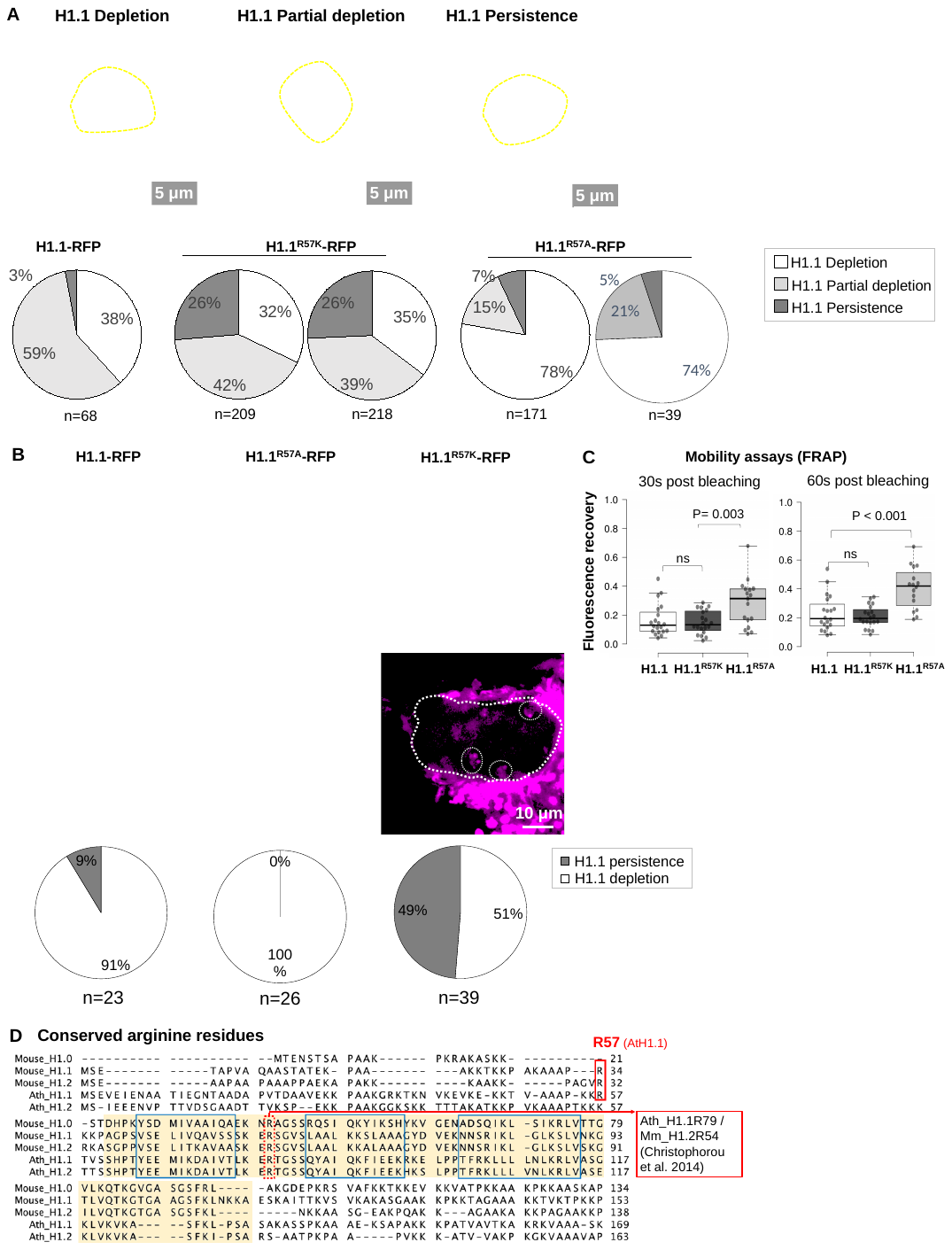

**Figure EV2 . H1.1 depletion in the SMC is controlled by the R57 residue (related to Figure 2)**

**Figure EV2 . H1.1 depletion in the SMC is controlled by the R57 residue (related to Figure 2)**

**(A)** Distribution patterns of H1.1 mutant variants showing either full depletion, partial depletion and persistence in the SMC (dashed lines) of ovule primordia stage 1-II/2-I at 5dpi (images, partial confocal projections) as used for scoring shown below for the H1.1-RFP control line and two independent lines expressing the mutant variants as indicated. n, number of primordia scored. **(B)** H1.1-RFP and H1.1^R57A^-RFP are evicted in male SMC (pollen mother cells), but H1.1^R57K^-RFP shows residual signal. Images are of premeiotic sporangia from flowers at 3dpi; pie charts quantify the observations. n, number of sporangia **(C)** Fluorescence revovery rate from FRAP experiments shown Figure 2, at 30s and 60s post bleaching. P value, Mann Whitney U test. **(D)** Alignment of selected Arabidopsis and Mouse variants as those in Figure EV1 (cropped around the globular domain, yellow, with parts of the N- and C-tails) showing the conservation of arginine residues (red boxes). The R57 residue from AtH1.1 studied in this work is positioned in the N-tail just before the globular domain. It is conserved in the Mouse H1.1 and H1.2 variants but not in the Arabidopsis H1.2 variant. The arginine shown by Christophorou and colleagues to be citrullinated is R54 in the Mouse H1.2 variant corresponds to R79 in AtH1.1 and AtH1.2. Data used for graphs and charts in **Dataset EV 2.**

**Figure EV3. The AIH citrullinase mediates H1.1 depletion in SMC (related to Figure 3)**

**(A)** Selected view of *AIH* expression generated by the ePlant Browser (bar.utoronto.ca/eplant/) showing middle-to-strong expression in young flower buds (box). **(B)** Schematic representation of the five splice variants of *AIH* (TAIR resource, arabidopsis.org) and the position of the probe used for RNA *in situ* hybridization shown below. Top panel: cross section through carpels, Bottom panel: detailed view of ovule primordia from the corresponding carpels at stages as indicated. **(C)** Position and sequence of the amiRNA used for downregulating *AIH* . **(D-E)** Representative images and scoring showing the effect of *AIH* downregulation using an amiRNA (D) or inhibition using Cl-amidine (E). Top panels: images of mock or Dex (D), or mock or Cl-amidine (E) treated ovule primordia at 5dpi expressing H1.1-GFP under its native promoter (She *et al*, 2013) and the inducible *amiR[AIH]* (D) as described in the main text. Dashed line: SMC. Pie charts: replicate scoring of depletion / persistence pattern of H1.1-GFP in the SMC, in three independent lines. n, number of ovule primordia scored, P values from Fisher exact tests. Data used for the charts in **Dataset EV 3.**

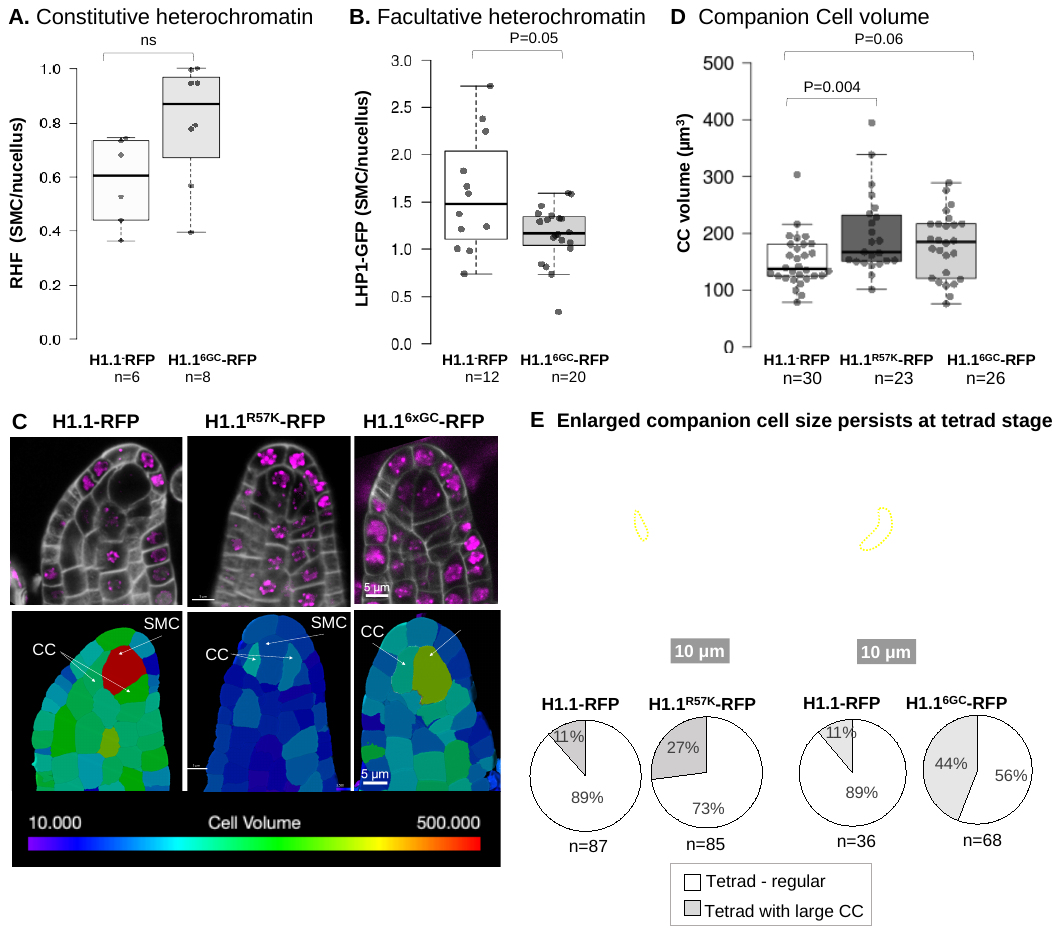

**Figure EV4. Effect of H1.1 persistence in the SMC on chromatin, SMC maturation and meiosis (related to Figure 4).**

**(A-B)** Relative Heterochromatin Fraction (RHF, A) and LHP1-GFP levels (B) in the SMC relative to surrounding nucellar cells, in the H1.1-RFP control line and H1.1^6xGC^-RFP mutant line. **(C)** Representative images and corresponding cell-based segmentation (below) of ovule primordia at 5dpi expressing the control of mutant H1.1 variants as indicated, to measure the SMC and CC volumes as plotted in Figure 4 and Panel D here. **(D)** Volume of the companion cells (CC) of ovule primordia at 5dpi expressing the control or mutant H1.1 variants as indicated. **(E)** Representative images of ovules at the tetrad stage showing a tetrad (white dotted line) and a neighboring, narrow or enlarged CC (left and right, respectively, yellow dotted line). Pie charts showing scoring of the respective classes in ovule primordia at 6dpi, induced for the expression of the control or mutant H1.1 variants as indicated. n, number of ovule primordia scored. P values, Mann Whitney U test. Data used for the graphs and charts in **Dataset EV 4**

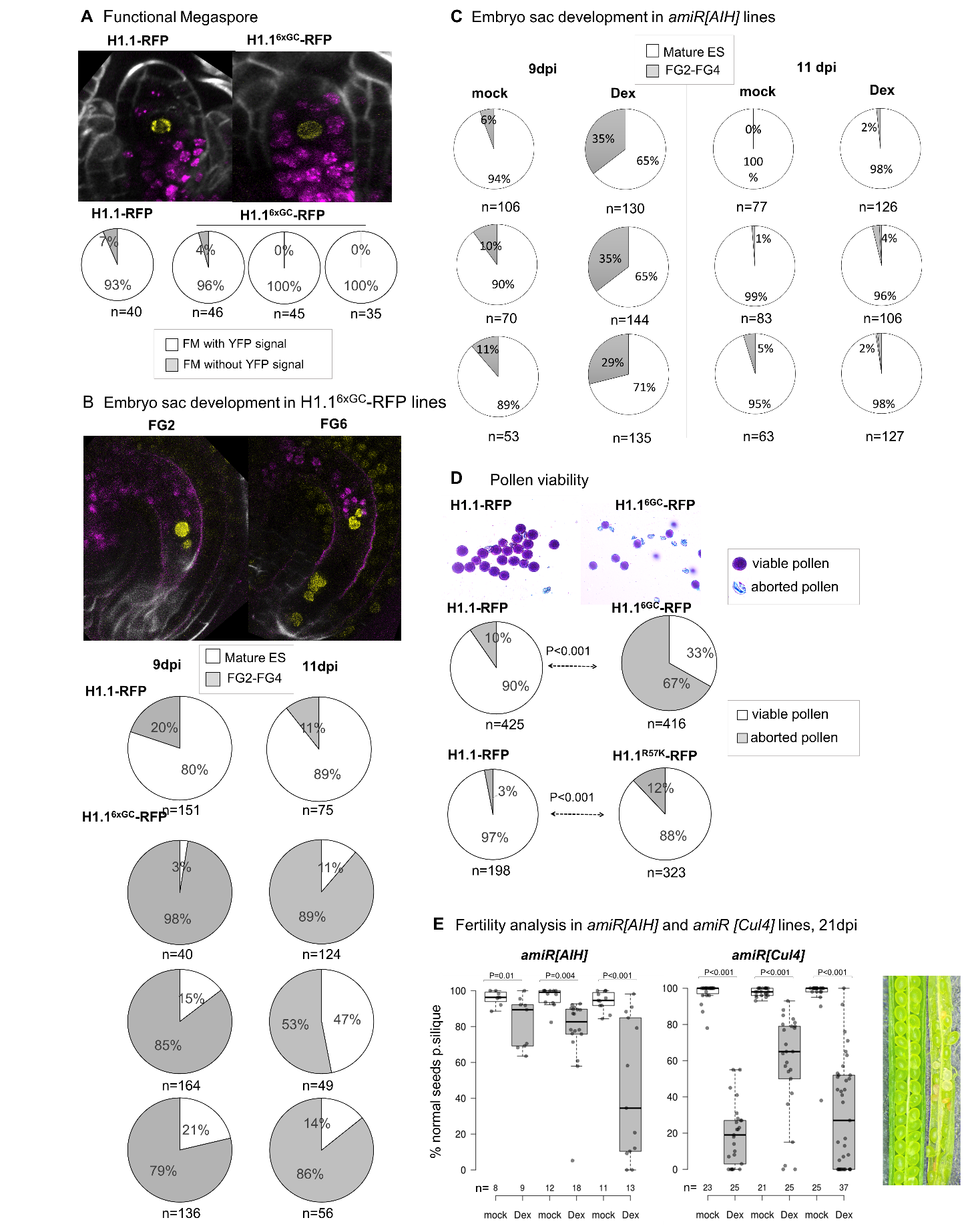

**Figure EV5. Impact of H1.1 persistence in the SMC on embryo sac development and fertility (related to Figure 5).**

**Figure EV5. Impact of H1.1 persistence in the SMC on embryo sac development and fertility (related to Figure 5).**

**(A)** Representative images and scoring of ovule primordia at 7dpi expressing *AKV::H2B-YFP* in the functional megaspore in the control or H1.1^6GC^-RFP line (3 independent lines). **(B)** Representative images and scoring of ovules at 9dpi and 11dpi (two days after emasculation at 9dpi) expressing *AKV::H2B-YFP* in the embryo sac (stages FG2 and FG6 are shown) in the control line and scoring of these classes in the control and H1.1^6GC^-RFP lines (3 independent lines). **(C)** Scoring of FG2-FG4 and FG6 embryo sacs identified by clearing, in ovules at 9dpi and 11dpi (two days after emasculation at 9dpi) following the induction of *amiR[AIH]* **(D)** Assessment of pollen viability by Alexander staining in the control and mutant lines as indicated, by scoring in anthers at 9dpi. P value, Fisher exact test. **(E)** Fertility analysis by quantifying the % of normal (green, plump) seeds per silique at 21dpi after mock or dex treatment, in mutant lines as indicated, in three independent lines. n, number of siliques scored. P values from a Mann-Whitney U test. Data used for graphs and charts in **Dataset EV 5**

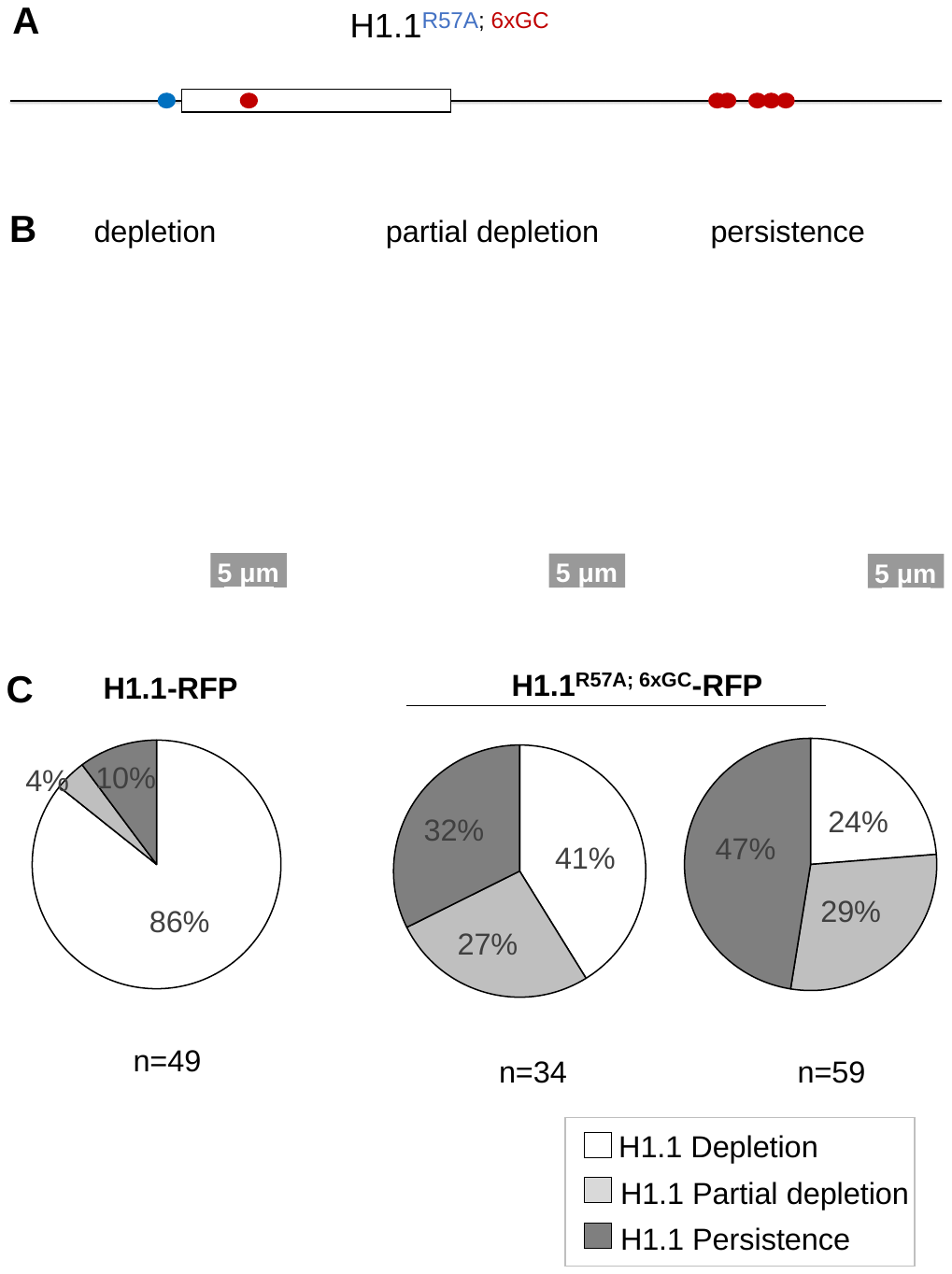

**Figure EV6. An H1.1 variant combining the R57A and K89R mutations shows resistance to degradation**

**(A)** Schematic representation of the H1.1^R57A;6xGC^ mutant variant showing the mutated R57A residue (blue) and the 6 K-to-R substitutions (red) among which only K89 in the globular domain (box). **(B)** Representative images of the depletion, partial depletion and persistence phenotype as scored in (C). Note the diffused pattern in the ‘partial’ category indicating increased dissociation from heterochromatin suggested to be a result of R57A. **(C)** Scoring in one control line and two independent double mutant lines as indicated. n, number of ovule primordia scored at 5dpi. Data used for charts in **Dataset EV 6**
